# Vacuole-specific lipid release for tracking intracellular lipid metabolism and transport in *Saccharomyces cerevisiae*

**DOI:** 10.1101/2021.05.04.442581

**Authors:** Vladimir Girik, Suihan Feng, Hanaa Hariri, W Mike Henne, Howard Riezman

**Author notes:** Present address: Unit of Chemical Biology and Lipid Metabolism, Center for Microbes, Development and Health (CMDH), Institut Pasteur of Shanghai, Chinese Academy of Sciences, Shanghai, China. Present address: Department of Biological Sciences, Wayne State University, Detroit, MI, USA. These authors contributed equally (Vladimir Girik, Suihan Feng).

## Abstract

Lipid metabolism is spatiotemporally regulated within cells, yet intervention into lipid functions at subcellular resolution remains difficult. Here we report a method that enables site-specific release of sphingolipids and cholesterol inside the vacuole in *Saccharomyces cerevisiae*. Using this approach, we monitored real-time sphingolipid metabolic flux out of the vacuole by mass spectrometry and found that the ER-vacuole tethering protein Mdm1 facilitated the metabolism of sphingoid bases into ceramides. In addition, we showed that cholesterol, once delivered into yeast using our method, could restore cell proliferation induced by ergosterol deprivation, overcoming the previously described sterol-uptake barrier under aerobic conditions. Together, these data define a new way to study intracellular lipid metabolism and transport from the vacuole in yeast.

## INTRODUCTION

Lipids are heterogeneously distributed in eukaryotic cells with their composition varying among the specific subcellular compartments (Harayama and Riezman 2018). This heterogeneity is achieved even though most lipid species are first synthesized in one organelle and transported to other organelles for use or further modification. Although most enzymes responsible for lipid metabolism have been identified, how lipids are transported within cells is much less known and many lipid transportation routes are still waiting to be established. Importantly, new methods are needed to be able to quantify the flux of lipids between intracellular compartments.

In recent years, a number of lipid transfer proteins (LTP) have been found to facilitate non-vesicular lipid transport through contact sites between organelles (Wong, Gatta et al. 2019). In many cases, structural information and *in vitro* lipid transfer assays need to be established to validate the roles of LTPs in lipid transport. This can be a challenging task because many LTPs are transmembrane proteins and lack detailed structural information. Mass spectrometry (MS)-based lipidomics techniques can contribute to our understanding of transport by providing quantitative analysis of many chemically diverse lipid species. Designed to carry out lipidome-wide scans, it has been particularly useful when combined with genetic perturbation in yeast or mammalian cells (Ejsing, Sampaio et al. 2009, Guan, Souza et al. 2009, da Silveira Dos Santos, Riezman et al. 2014, Jimenez-Rojo, Leonetti et al. 2020). Nevertheless, the results of gene deletion do not always translate into lipid profile changes. For example, simply removing one LTP may not generate changes in lipid profiles if a lipid transportation route is maintained by several proteins with overlapping functions, as seems to be often the case. Clearly, it is desirable to develop new approaches which overcome gene redundancy issues and do not rely on *in vitro* protein-lipid reconstruction.

Previously, we have developed a series of coumarin-based photo-cleavable (caged) lipid probes targeted to mitochondria and lysosomes, respectively (Feng, Harayama et al. 2018, Feng, Harayama et al. 2019). Upon illumination, these probes quickly decomposed, thereby releasing the corresponding native lipid molecules inside the targeted organelle. Using isotope-labelled sphingosine as a lipid precursor, we detected its metabolic products by mass spectrometry with high sensitivity, and showed that sphingolipid metabolic patterns depend highly on subcellular localization (Feng, Harayama et al. 2019). While this strategy is firmly established in mammalian cells, it was unclear whether the same concept is applicable to yeast, an extensively used model organism for studying lipid homeostasis owing to its simplicity, ease of genetic manipulation and similar organization to metazoans on cellular and subcellular levels.

To explore their use in yeast, we applied mitochondrial-targeted caged probes onto *Saccharomyces cerevisiae (S. cerevisiae*) and found surprisingly that these probes accumulated inside the vacuole instead of mitochondria. This unexpected finding led us to synthesize a series of caged lipids, including sphinganine, phytosphingosine (PHS) and cholesterol, which were applied to investigate vacuole-derived lipid metabolism (Figure 1A). Using mass spectrometry as a readout, we characterized the metabolic products of vacuolar sphingoid bases, explored potential effectors on the sphingolipid metabolic pathway, and provided clear evidence that an ER anchored protein which also contacts the vacuole surface, Mdm1, facilitated sphingolipid turnover. Additionally, we successfully delivered cholesterol into *S. cerevisiae* using the same strategy. We showed that cholesterol can restore cell proliferation induced by ergosterol deprivation, and that deletion of NCR1, the orthologue of NPC1, which is important for cholesterol transport in mammals (Zhang, Ren et al. 2004, Gong, Qian et al. 2016), did not strongly affect vacuole-derived cholesterol utilisation in yeast cells. Collectively, our approach, which combines synthesis, imaging, lipidomics and genetic modification, defined a new way of studying intracellular lipid transport and metabolism in *S. cerevisiae*.

**Figure 1.**
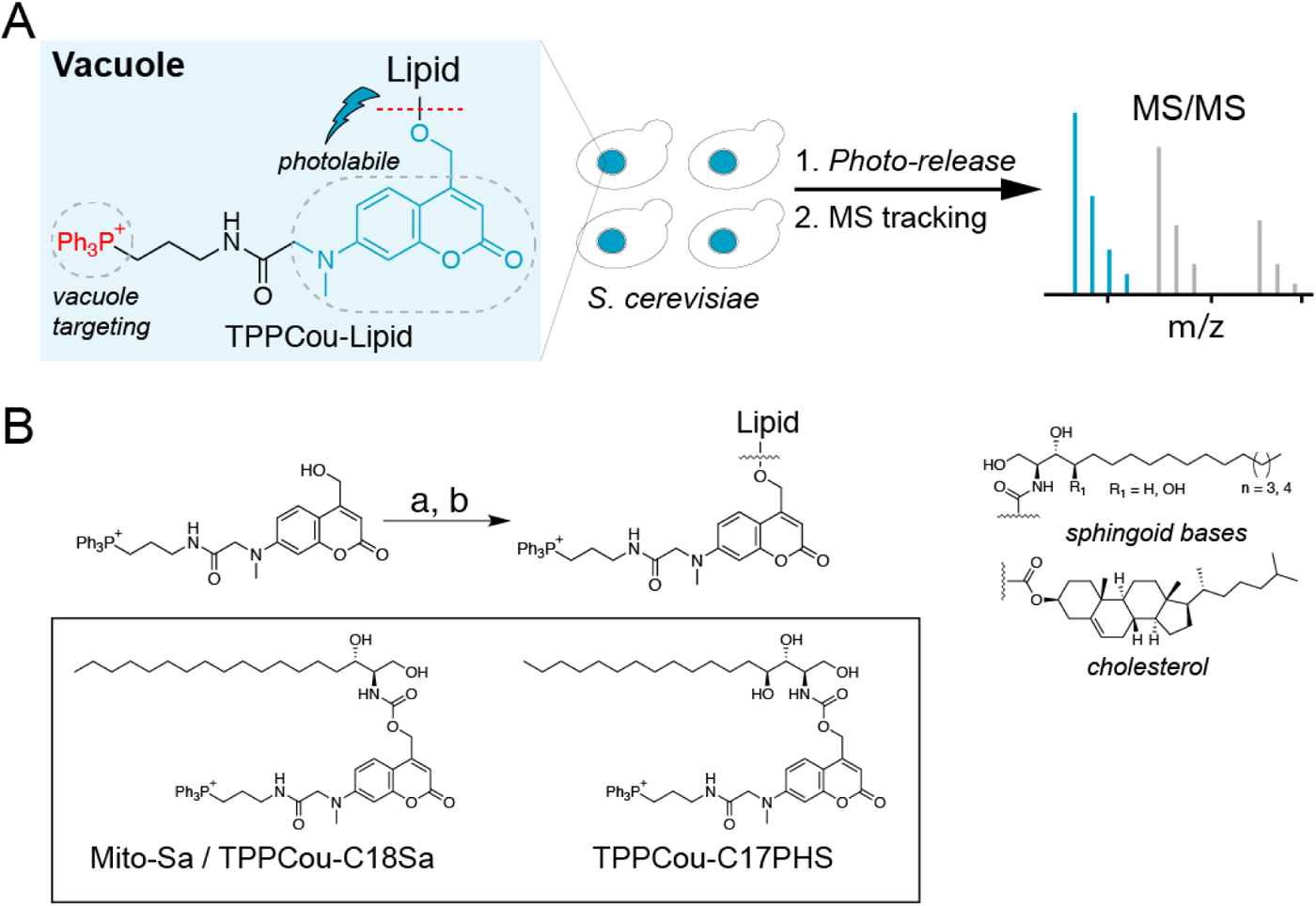
(A) Schematic illustration of MS-based metabolic tracking of vacuole-targeted photocleavable (TPPCou) lipid probes. (B) Synthesis of TPPCou-caged lipids and chemical structures of TPPCou caged sphingoid bases. For long chain sphingoid bases (LCBs), *i*) Bis-(4-nitrophenyl)carbonate, DIPEA, DMF, r.t., 3 h; LCB, 60 °C 3 h, 40-60%. For TPPCou-Chol, cholesteryl chloroformate, DCM, DMAP, 18h, 60 %.

## RESULTS

### TPPCou-caged sphingoid bases accumulate in the yeast vacuole

The triphenylphosphonium (TPP) cation is a well-known mitochondrial targeting motif and has been used to deliver a wide range of small molecules into the mitochondrial matrix in mammalian cells (Murphy 2008). In the past, we have used TPP-modified coumarin as a photo-cleavable protecting group to prepare caged sphingolipids targeting to the mitochondria in mammalian cells. Unlike in Hela cells, TPP-coupling strategy has not been widely used in yeast, yet several studies in the pathogenic yeast *Candida albicans* showed that TPP-conjugated antifungal compounds inhibited multidrug resistance efflux pumps (Chang, Liu et al. 2018) and interfered with mitochondrial functions (Wang, Liu et al. 2021). In another study (Severin, Severina et al. 2010) TPP coupled to a fatty-acid anion stimulated respiration and thus was presumably targeted to mitochondria in *S.cerevisiae,* but no visual localization studies supporting mitochondrial localization were provided. Therefore, we examined the localization of TPP-coumarin caged compounds, which are easily seen by fluorescence microscopy, because the coumarin is also a fluorophore in addition to being a photo-cleavable protecting group. Since it was unclear whether these caged probes would accumulate in yeast mitochondria, we incubated the caged sphinganine (Mito-Sa, Supplementary Figure 1A) in *S. cerevisiae* expressing a fluorescent mitochondrial marker, Mdh1-mCherry. To our surprise, while we recorded a strong fluorescence signal in the coumarin channel, the staining pattern did not overlap with the Mdh1-mCherry signal, but rather looked like the vacuole (Supplementary Figure 1B). This unexpected finding prompted us to examine co-localization with a vacuole marker, Vph1-mCherry, which consistently overlapped well with the coumarin fluorescence (Supplementary Figure 1C). Since Mito-Sa localized to the mitochondria in Hela cells, but to the vacuole in yeast cells, we renamed it **TPPCou-C18Sa** in order to not cause any confusion.

Since phytosphingosine (PHS) is the major form of long chain sphingoid bases (LCBs) in yeast (Haak, Gable et al. 1997), we synthesized and purified TPPCou-C17PHS, to distinguish it from the native C18PHS, (Figure 1B) and investigated its localization in *S. cerevisiae*. Using fluorescence microscopy, we have found, consistently, that TPPCou-C17PHS accumulates inside the vacuole, but not the mitochondria (Figure 2 A-D). To learn which factors affect the vacuole staining, we treated the cells with diethylaminocoumarin caged PHS (Cou-PHS) without the TPP cation and TPPCou alone, respectively (Supplementary Figure 2 A-C), but both failed to generate any meaningful fluorescence signals, indicating that both the TPP cation and a lipid chain are compulsory for effective transport of caged probes into the vacuole. It is possible that TPPCou was pumped out by yeast efflux pumps (Ernst, Klemm et al. 2005) since the TPP cation could be delivered into yeast cell only by applying pulsed electric fields (Stirke, Zimkus et al. 2014). Due to the dependence on the sphingoid base, we postulated that proteins involved in long-chain base uptake may play a role in the internalization of TPPCou-C17PHS. Indeed, we found a significant decrease of fluorescence signals in the absence of fatty acyl-CoA synthetases (*faa1*, *faa4*) and/or transporter (*fat1*), which have been shown to affect uptake of exogenous sphingoid bases (Narita, Naganuma et al. 2016) (Figure 2E). Next, we treated the cells with TPPCou-C17PHS in the presence of CCCP and bafilomycin A1, respectively. Either disrupting the proton gradient with CCCP or blocking vacuolar V-ATPases with bafilomycin significantly reduced the fluorescence intensities of TPPCou-C17PHS, suggesting that the vacuole staining is dependent on the pH gradient across the vacuolar membrane (Figure 2F). Furthermore, we investigated whether the uptake of TPPCou-C17PHS was via endocytosis. Using cells that lack *END3*, essential for endocytosis (Benedetti, Raths et al. 1994), we found that the fluorescence intensities from *end3* cells are even slightly higher than the ones from the wildtype (*WT*) cells (Figure 2G), suggesting that the transport does not rely on endocytosis. Notably, the average fluorescence intensities in both conditions (Figure 2G) are relatively lower compared to other experiments (Figure 2E, F), likely caused by the differences among the yeast strain backgrounds. Together with the imaging data of TPPCou-C18Sa (Supplementary Figure 1), we showed that the TPPCou caged lipids accumulated inside the yeast vacuole, and that elements crucial for the import and localization include the proton gradient across the vacuolar membrane and proteins required for sphingoid base and fatty acid import.

**Figure 2.**
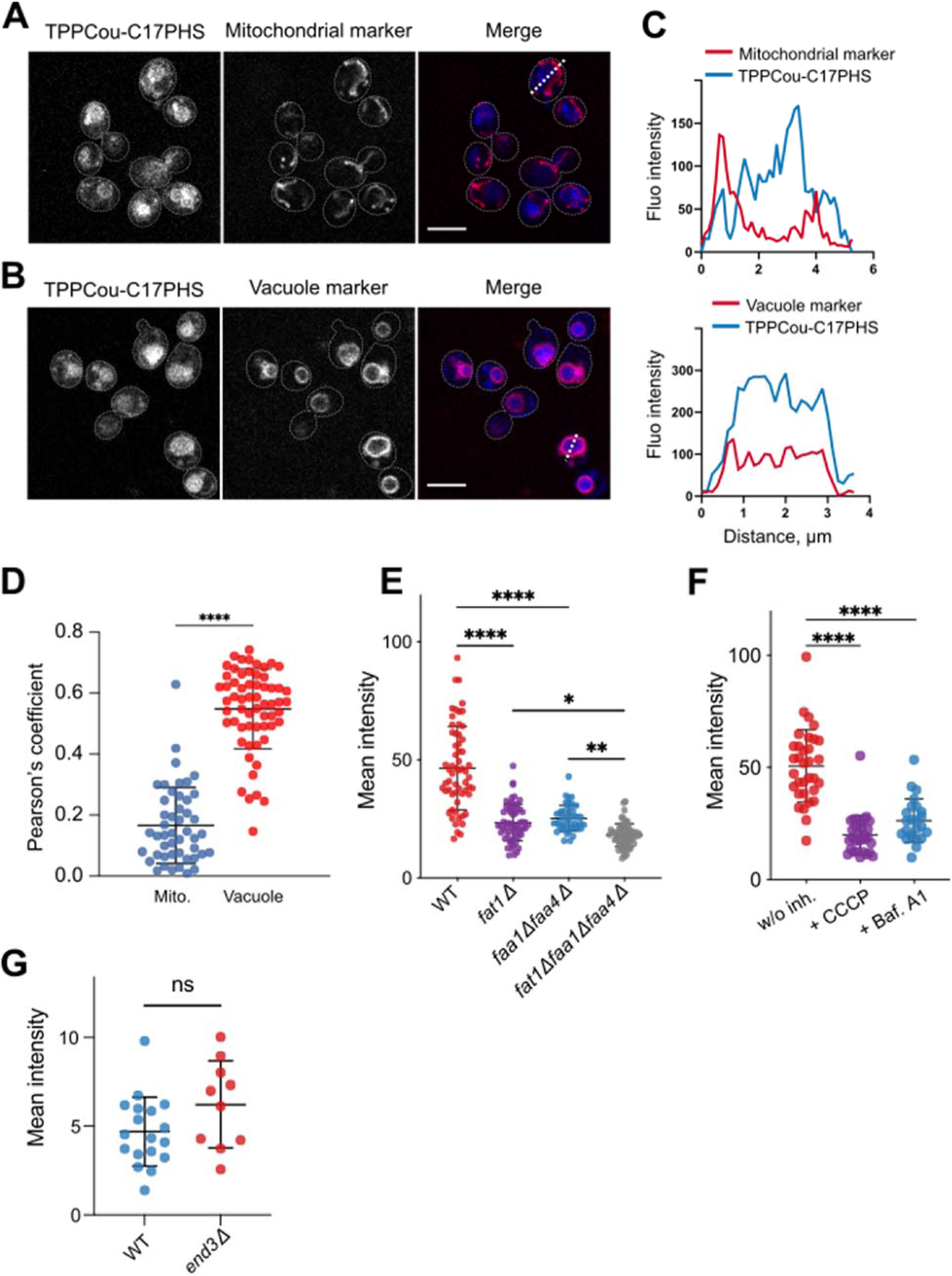
TPPCou-C17PHS accumulated in the yeast vacuole. (A) and (B) Representative fluorescence images of yeast cells stably expressing mCherry-tagged (red) mitochondrial marker protein Mdh1 (A) or vacuolar protein Vph1 (B) stained with 10 µM TPPCou-C17PHS (blue). Bars, 5 µm (C) Intensity plots of white dotted lines shown on (A) and (B) for the indicated fluorescent channels. (D) Quantification of colocalization of TPPCou-C17PHS with a mitochondrial marker (Mdh1-mCherry) and vacuole marker (Vph1-mCherry), respectively. (E, F, G) Quantification of mean intracellular fluorescence intensity of TPPCou-C17PHS. Cells were treated with 20 μM TPPCou-C17PHS for 15 min in the presence of indicated conditions. *p<0.05, **p<0.01, ****p<0.0001, ns., not significant, student’s *t*-test.

### Metabolism of vacuole-released sphingoid bases is less dependent on LCB4

In *S. cerevisiae*, the levels of long chain sphingoid bases (LCB) are tightly regulated by coordinated action of metabolic enzymes (Breslow 2013). We have previously reported that Lcb4 kinase is essential for incorporating exogenous LCBs into complex sphingolipids (Funato, Lombardi et al. 2003), and its activity accounts for 95% of long-chain base kinase activity (Nagiec, Skrzypek et al. 1998). Lcb4 has been localized to the Golgi, late endosomes (Hait, Fujita et al. 2002), but more direct localizations without protein tagging found it on the cortical ER juxtaposed to the plasma membrane (Iwaki, Sano et al. 2007). Given the localization of Lcb4 outside of the vacuole, it is therefore interesting to know whether the conversion of vacuole-released sphingoid bases into dihydroceramides is also subject to regulation by Lcb4. To this end, we prepared a caged, deuterated C18 sphinganine probe (TPPCou-Sa-D7, Figure 3A) and analyzed its metabolic products after photo-releasing in wildtype (WT) and mutant cells lacking LCB4 (*Δlcb4*), respectively. In parallel, we used exogenously added C18 sphinganine (Sa-D7) as a control (Figure 3B). The downstream metabolites of the deuterated sphinganine include phytoceramides (PHC) and inositol phosphorylceramide (IPC), which can be detected by mass spectrometry (Figure 3C). Indeed, we observed that conversion of sphingoid base into PHC and IPC was drastically reduced in *Δlcb4* cells (Figure 3D, E), similar as our previous study (Funato, Lombardi et al. 2003). Decreased PHC and IPC synthesis was also found in the vacuole-released Sa-D7 experiments, but to a much lesser extent (Figure 3F, G). These results suggest that a large part of the Lcb4-dependence for complex sphingolipid production from exogenous sphingoid bases may be related to cellular uptake. However, this cannot be the entire dependence because some dependence was also seen *in vitro* after reconstitution (Funato, Lombardi et al. 2003). In contrast, our data here show that a large part of the recycling pathway of sphingoid bases from the vacuole into complex sphingolipids does not depend on LCB4, consistent with the previously described Lcb4 localization in proximity to the plasma membrane (Iwaki, Sano et al. 2007).

**Figure 3.**
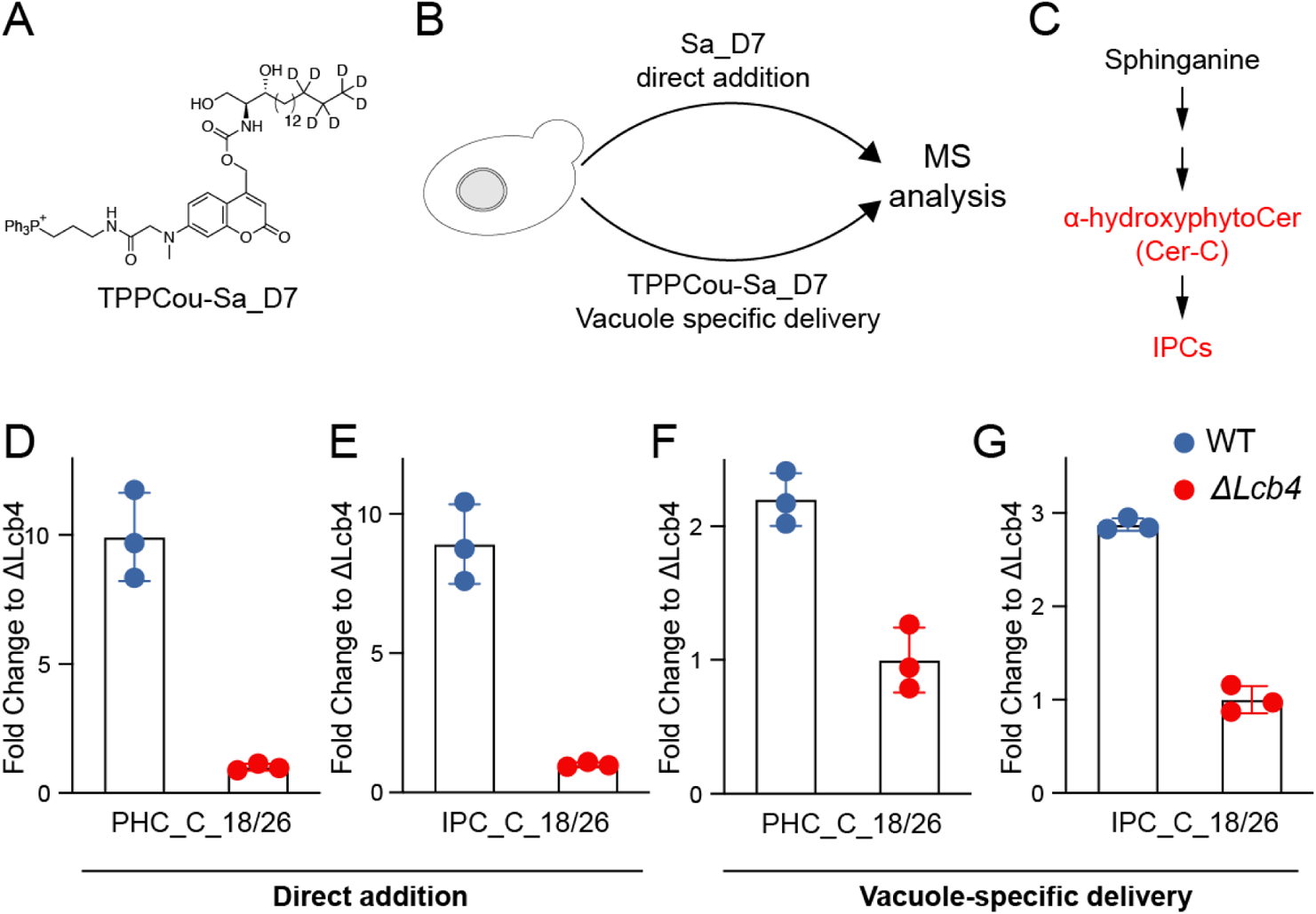
Comparison of D7-sphinganine conversion to phytoceramide and inositolphosphoceramide. (A) Schematic illustration of the experimental design. (B) Simplified metabolic routes of sphinganine. (C-F) Quantification of PHC_C_18/26 and IPC_C_18/26 using two D7-sphinganine delivery methods as indicated. Data represents three biological replicates. Error bars represent SD.

### Metabolism of vacuole-released C17PHS

The lysosome/vacuole serves as a primary site to break down complex lipid molecules into basic building blocks, which are later transported to other sites such as endoplasmic reticulum (ER) for lipid synthesis (Li and Kane 2009). These processes of storage and recycling are of fundamental importance and have been linked to a number of diseases, but many aspects, particularly on the sphingolipid transport, remain elusive (Platt 2014). Previously, we have applied the local uncaging methods in Hela cells to show that sphingolipid metabolism and turnover have distinct properties depending on the subcellular localization of the release (Feng, Harayama et al. 2019), though the mechanism of lipid transport was not defined. Here we applied the same principle to examine vacuole-released sphingolipid metabolism, with a particular focus on lipid transport.

The vacuole maintains a low pH environment and hosts a range of enzymes to disassemble lipid molecules. To assess the stability of caged probes after import into the vacuole, cells were treated with TPPCou-C17PHS, collected before or after UV illumination, and analyzed by mass spectrometry (Figure 4B). We could detect some C17PHS even without illumination, likely due to undesired enzymatic activities in the vacuole. Nevertheless, the majority remained intact, judging from the UV light-released C17PHS, which is about four times higher than the control. After learning that the majority of the caged probe was intact, we analyzed its metabolic products after uncaging and compared to the ones delivered by extracellular addition of C17PHS. Phytosphingosine can be transported from the vacuole to the nuclear ER where it is converted into various ceramides and other complex sphingolipids (Figure 4A, Supplementary Figure 3). Our analysis showed that exogenously added C17PHS was predominantly converted into Phytoceramide C (PHC-C_C43). In photo-released cells, however, a significantly higher proportion of C17PHS was used for the synthesis of Phytoceramide B (PHC-B_C_43) and shorter-chain ceramides (Figure 4C). Although the amount of C17PHS being delivered into intracellular space was different under the two experimental conditions, these results clearly revealed subcellular dependence of sphingolipid metabolism, in accordance with our previous findings in mammalian cells.

**Figure 4.**
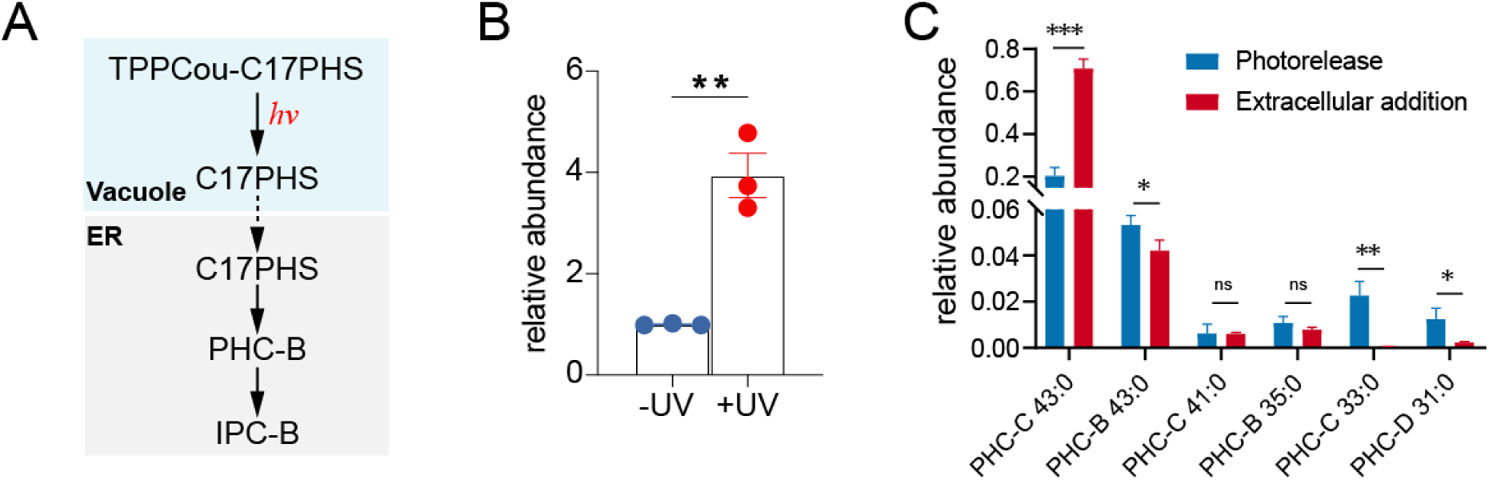
Metabolism of vacuole-released C17PHS. (A) Metabolic scheme of C17PHS after the vacuole specific-uncaging. After transport from the vacuole to the ER, PHS is converted into Phytoceramide (PHC-B) which is transported further to the Golgi, where PHC-B is metabolized into Inositol Phosphorylceramide (IPC-B). (B) Analysis of C17PHS levels before (-UV) and after(+UV) UV uncaging in cells labeled with 5 µM TPPCou-C17PHS. C) Comparison of ceramide species derived from TPPCou-C17PHS (after photo-release) or extracellularly added C17PHS. (E-H) Lipid analysis of sphingolipids derived from vacuole-released C17PHS after UV uncaging in cells overexpressing empty vector, MDM1 (E-F) or NVJ1 (G-H). Data represents the average of three independent experiments. Error bars represent SEM. *p<0.05, **p<0.01, ***p<0.001, ns, not significant, students’ *t*-test.

### MDM1 facilitated sphingolipid turnover

Lipids can be transported within cells by two types of pathways, vesicular and non-vesicular, but for most lipids it is thought that majority of traffic is carried out by non-vesicular trafficking, most likely involving lipid transfer proteins and membrane contact sites (MCS), areas where membranes from two organelles are found in close apposition (Prinz, Toulmay et al. 2020). Nucleus-vacuole junctions (NVJs) are an example of membrane contact sites between the vacuole and the nuclear ER wrapped around the nucleus forming a contiguous membrane (Pan, Roberts et al. 2000). A number of imaging-based studies indicated the universal presence of MCS, but functional studies are still lagging behind (Huang, Jiang et al. 2020). In the recent years, we have identified MDM1 as a tethering protein which localized to the ER but forms contacts with the vacuole and lipid droplets in yeast (Hariri, Rogers et al. 2018, Hariri, Speer et al. 2019). MDM1 is highly conserved in metazoans and plays essential roles in regulating lipid homeostasis (Datta, Liu et al. 2019, Ugrankar, Bowerman et al. 2019). We have found that mutations in yeast MDM1 perturb sphingolipid metabolism, but whether this tether is involved in inter-organelle lipid trafficking remained elusive (Henne, Zhu et al. 2015). In support of a potential lipid trafficking model, recent structural predictions using the machine-learning algorithm Alphafold2 indicate that MDM1 and its orthologs contain a putative novel lipid transport region formed from two of its domains, the PXA and PXC, which fold intra-molecularly into a bi-domain module with a large hydrophobic cavity (Paul, Weeratunga, et al. 2022). Motivated by these observations, we next examined whether MDM1 may influence ER-vacuole lipid trafficking.

We used the vacuole-specific uncaging method to examine whether sphingolipid turnover is influenced by MDM1 over time. Accordingly, we developed a pipeline in which C17PHS was first released inside the vacuole, and its metabolic products were measured by mass spectrometry at different time points. PHC-B_C43 and IPC-B_C43 are two of the major metabolic products derived from C17PHS and gave lowest signal-to-noise ratios, and hence were selected in our analysis. Our data showed that the accumulation of both PHC-B_C43 and IPC-B_C43 were significantly higher in cells in which MDM1 was overexpressed (Figure 5A, B). However, when we overexpressed NVJ1, another established nuclear ER-vacuole junction (NVJ) tether (Pan, Roberts et al. 2000), we did not observe any significant difference when compared to the control cells (Figure 5C, D). This striking contrast between MDM1 and NVJ1 overexpression, which both promote increased MCS formation, indicates that MDM1 promoted the turnover of vacuolar sphingolipids by additional means than its role in MCS formation.

**Figure 5.**
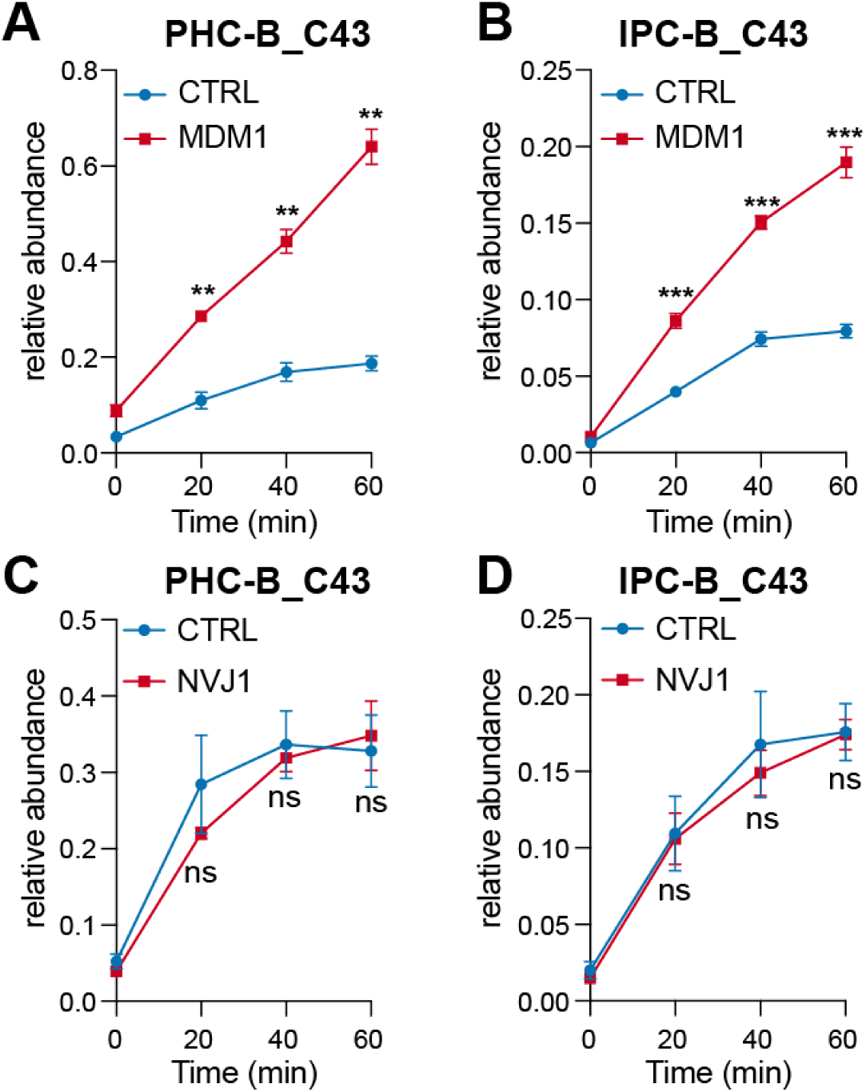
MDM1 facilitated sphingolipid turnover. Lipid analysis of sphingolipids derived from vacuole-released C17PHS after UV uncaging in cells overexpressing empty vector, MDM1 (A, B) or NVJ1 (C, D). Data represents the average of three independent experiments. Error bars represent SEM. *p<0.05, **p<0.01, ***p<0.001, ns, not significant, students’ *t*-test.

Intriguingly, when we attempted to use MDM1 or NVJ1 knockout strains in the ‘pulse-chase’ experiments, we detected high levels and variable quantities of endogenous C17 sphingolipids. Since those sphingolipids share the same chemical structures with the ones converted from the C17PHS after uncaging, it was difficult to deduce any conclusion from those experiments. The reason for the appearance of odd-chain sphingolipids in these mutants is unknown. Since neither MDM1 nor NVJ1 is essential for cell growth, cells do not seem to rely on a single component/pathway to transport lipids between the vacuole and the ER. The functional redundancy and/or quick metabolic adaptation could also mean that overexpression rather than gene deletion is more suitable to capture differences in the metabolic tracking studies using our method. In addition, our lipidomics analysis indicated that major lipid profiles were not significantly shifted after the photo-release (Supplementary Figure 4), most likely because the LCBs delivered to the cells are tracers and only comprise a very small portion of the endogenous lipid pool.

### Cholesterol delivered to yeast using TPPCou as cargo

Cholesterol is a fundamentally important molecule to human health, yet its physiological roles are still not clearly defined and some are highly debated (Goldstein and Brown 2015). This is partially because cholesterol is so abundant and essential in mammalian cells that any experimental means for reducing cholesterol levels faces the risk of jeopardizing numerous cellular functions. In contrast, wildtype yeast does not produce cholesterol and use another sterol, ergosterol, for maintaining cellular activities. Having engineered a yeast strain that effectively produces cholesterol, we demonstrated previously that functions of ergosterol can be partially replaced by cholesterol, which makes yeast an attractive model organism for studying cholesterol transport and metabolism (Souza, Schwabe et al. 2011). However, while mammalian cells import cholesterol through receptor mediated endocytosis, *S. cerevisiae* does not take up sterols under aerobic conditions (Keesler, Casey et al. 1992).

To gain insights into cholesterol transport, we prepared TPPCou caged cholesterol (TPPCou-Chol, Figure 6A) and inspected whether it can be effectively delivered to the vacuole in wild type yeast cells. As hoped for, the probe was successfully delivered into the vacuole, like the TPPCou-C17PHS. In addition, we measured the cholesterol uptake under various conditions and found that cholesterol was indeed delivered to the cells using TPPCou-Chol, although there was only minimal additional effect due to UV illumination (Figure 6D). These results suggest that most TPPCou-Chol was cleaved by enzymes prior to UV exposure, which also partially explains why the fluorescence signals from TPPCou-Chol were weaker than the ones from TPPCou caged LCBs (Figure 6B, C). Despite lacking optical control on the TPPCou-Chol, the data marked the successful vacuole-targeted delivery of cholesterol under aerobic conditions, enabling us to further investigate cholesterol transport in *S. cerevisiae*. First, we blocked endogenous ergosterol biosynthesis using fenpropimorph (Marcireau, Guilloton et al. 1990), then treated cells with TPPCou-Chol in increasing amounts, and monitored cell growth. The growth curves indicated that TPPCou-Chol effectively rescued the deficiency of ergosterol in a dose-dependent manner (Figure 6E), while free cholesterol treated cells failed to alleviate any cell growth suppressed by fenpropimorph (Figure 6F). Using fenpropimorph, we also performed another set of experiments on NCR1 knockout cells. We observed similar growth curves positively correlated with the amount of TPPCou-Chol, but did not see any significant difference between wild type (*WT*) and NCR1 knock cells (*Δncr1*) (Figure 6G, H). Our results suggest that NCR1 is not essential for transporting cholesterol from the lysosomes/vacuoles, in agreement with suggestions from previous findings (Malathi, Higaki et al. 2004, Zhang, Ren et al. 2004). Overall, our data showed that the vacuole-specific uncaging approach is not limited to sphingoid bases and has the potential for broader application to study lipid recycling. Furthermore, we have established a novel protocol to introduce sterols into aerobically grown yeast, overcoming the sterol exclusion mechanism, which should permit future studies on the function of sterols in cell biology.

**Figure 6.**
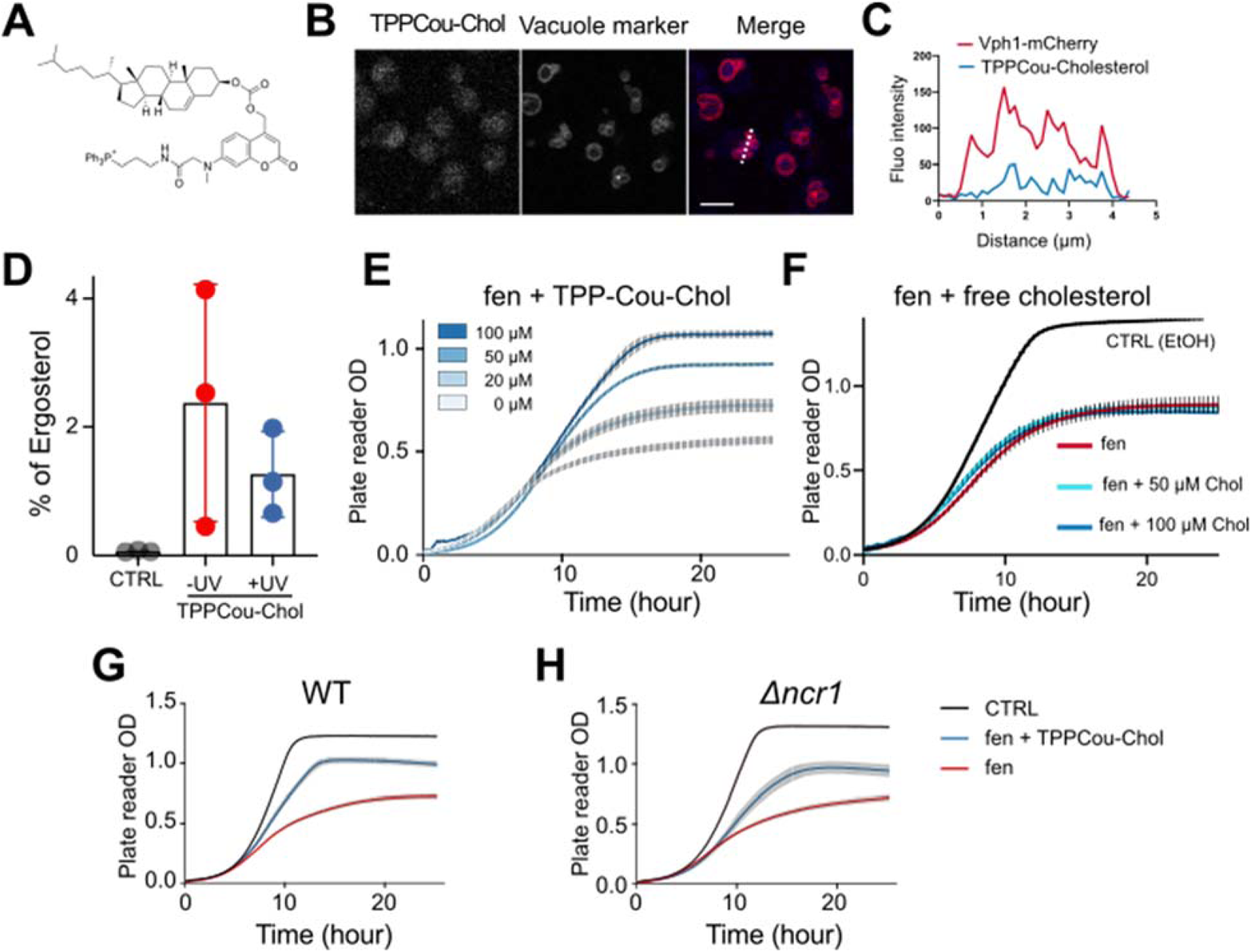
Vacuole-released cholesterol rescues growth of ergosterol-depleted yeast cells. (A) Chemical structure of TPPCou-Chol. (B) Representative confocal images of yeast cells stably expressing mCherry-tagged (red) vacuolar marker protein Vph1 stained with 50 µM TPPCou-Chol (blue) for 20 min at 30°C. Bars, 5 µm. (C) Intensity profile of the white dotted line in (B) in TPPCou-Cholesterol and Vph1-mCherry channel, respectively. (D) GC-MS analysis of vacuole-released cholesterol in yeast cells treated with 50 µM TPP-Cou-Choleserol or ethanol (CTRL) and subjected to UV uncaging (+UV). Cholesterol amounts are shown as percentage of total ergosterol levels. Data represents the average of three independent experiments. Error bars represent *SEM*. (E) Growth curves of cells treated with TPPCou-Chol in the presence of fenpropimorph (8 µM). (F) Growth curves of cells treated with free cholesterol in the presence of fenpropimorph (8 µM), or ethanol (CTRL) as indicated. (G, H) Growth curves of WT or *ncr1Δ* mutant cells. Cells were grown in presence of fenpropimorph (8 µM),TPPCou-Chol or control (ethanol). Cells were incubated in a plate reader and the growth was recorded by taking regular OD measurements at 30°C. Individual values represent the average of four biological replicates. Error bars represent SEM.

## DISCUSSION

In eukaryotic cells, distinct lipid distribution and metabolism are associated with subcellular compartments, but related tools to directly address these issues are very limited. Recently, we have developed a local uncaging strategy that offers direct experimental means to study organelle-specific lipid metabolism. Here, we introduced this concept in the yeast system and we have extended its application to intracellular lipid transport. Together with mass spectrometry, we provide quantitative information of sphingolipid metabolism, its dependence on sphingosine kinase, and unveiled the active role of MDM1 in facilitating the conversion of long chain sphingoid bases (LCBs) into ceramides. In addition, by releasing cholesterol in the vacuole, we showed that cholesterol can effectively rescue the deficiency of ergosterol, that cholesterol transport was not substantially hindered by the removal of NCR1 and that it is possible using this method to introduce sterols into aerobically grown yeast.

The fluorescence imaging data shows that TPPCou caged lipids accumulated in the vacuole, but it is still unclear why they are enriched inside vacuole instead of mitochondria. Triphenylphosphonium (TPP) cation is known as a mitochondrial targeting signal and has been extensively used in Hela cells, yet we did not find any targeting to mitochondria in yeast. Previously, anti-fungal TPP-coupled compounds could be targeted to mitochondria in the pathogenic yeast C*andida albicans* (Chang, Liu et al. 2018, Wang, Liu et al. 2021). Another study demonstrated that TPP-conjugated with a fatty acid stimulated yeast mitochondrial respiration (Severin, Severina et al. 2010) but the localization of the compound was not addressed and its effects on mitochondria could have been indirect. It has been known that cationic chemotherapeutic drugs are trapped in lysosomes, organelles analogous to the yeast vacuole (Kaufmann and Krise 2007). It is also known that the budding yeast express multiple H^+^-drug antiporters and at least one of which, Vba4, is found in the vacuolar membrane (Kawano-Kawada, Pongcharoen et al. 2016). The activity of H+-antiporters depends on the vacuolar pH gradient and experiments using CCCP and bafilomycin indicate that the pH gradient is essential to localize the TPP-Cou probes inside the vacuoles. Although CCCP is commonly used as an uncoupler of mitochondrial potential it has also been shown to equilibrate the yeast vacuolar pH with that of the extracellular medium (Padman, Bach et al. 2013). It is difficult to rule out the possibility that some of the TPPCou-based probes went into mitochondria, but the amount accumulated in mitochondria is under the detection threshold and thus must be limited. We explored the uptake mechanism of TPPCou caged lipids using knockout strains, and our results show that probe uptake involves the participation of acyl-CoA synthases but is independent of endocytosis. Since the vacuole is a recycling hub for breaking down complex lipids into building blocks, it is not a surprise that we observed partial hydrolysis of TPPCou-C17PHS prior to UV illumination, and that TPPCou-Chol was completely decomposed. This cleavage should have occurred in the vacuole, not before delivery there, because incubation with TPPCou alone did not lead to vacuolar labeling. On the other hand, the successful delivery of long chain sphingoid bases and cholesterol also suggests a possible broader application of using TPPCou as cargo to transport other lipid molecules to the vacuole in *S. cerevisiae*.

The major lipid trafficking routes are organized by non-vesicular lipid transport, particularly through membrane contact sites (MCS) with the involvement of a large group of lipid transfer proteins (LTP). Several imaging-based studies revealed the precise localization of these proteins, but functional studies often lag behind. Herein, we released phytosphingosine in vacuole and monitored its metabolic products over time, to establish a system which can directly measure the influence of LTPs on lipid movement and metabolism. Our results showed that overexpressing MDM1, an ER-vacuole tethering protein, facilitated formation of ceramide and IPC species, unlike another nuclear ER-vacuole junction protein, NVJ1. Since ceramide synthases are localized to the nuclear ER, active lipid transport from vacuole to the ER is a prerequisite before the metabolic conversion. We have previously shown that MDM1 mutants suppress cell survival in myriocin treated plating assays (Henne, Zhu et al. 2015), but details were lacking because suppressing sphingolipid biosynthesis by myriocin can have profound effects on numerous aspects of cellular activities. In our current study, we measured the real-time metabolic flux of sphingolipids originating from the vacuole, thus providing a direct link between MDM1 and the metabolic turnovers of sphingoid bases. Our recent studies (Hariri, Rogers et al. 2018, Hariri, Speer et al. 2019) also found that MDM1 directly interacts with fatty acids via its hydrophobic N-terminal region and promotes lipid droplet formation. It is unclear for now whether MDM1 directly binds long chain sphingoid bases, but even if the direct interaction occurs, it is likely that the mechanisms are somewhat different because the head group of sphingoid bases is much more polar and hydrophilic than fatty acids. However, it is notable that recent structural predictions using Alphafold2 suggest that MDM1 and its orthologs encode a putative LTP-like module composed of its PXA and PXC domains (Paul, Weeratunga, et al. 2022). An intriguing model is that this LTP-like module is capable of transporting lipids including C17PHS at inter-organelle contacts.

We also demonstrated that the same principle can be used to study cholesterol transport in yeast using growth as a functional readout, highlighting the flexibility and compatibility of our approach for studying local lipid metabolism. As cholesterol plays essential roles in maintaining numerous cellular functions, switching the system to *S. cerevisiae* should offer more freedom to modulate its levels and to introduce analogs.

In metabolic flux studies, our method should be able to scan multiple lipid transfer protein candidates without the requirement of detailed information on protein structures. Arguably, our design relies on biochemical conversion and hence enzymatic activities, but this requirement should be matched without difficulty thanks to the abundance of numerous lipid metabolic enzymes (Supplementary Figure 3). In future applications, we can use lipid molecules bearing a clickable motif and a diazirine which allows crosslinking to proteins in close proximity after exposing to an orthogonal UV light to photo-uncage, as has been previously demonstrated (Hoglinger, Nadler et al. 2017). In this case, after photo-uncaging, crosslinking followed by ‘click’ chemistry with a fluorophore should enable us to visualize lipid localization over time without relying on enzymatic activities, though it will lack the information of lipid species.

In conclusion, we presented here a novel technique of delivering lipid precursors specifically to the vacuole in *S. cerevisiae*, which enabled us to visualize and track lipid recycling from the vacuole. It has also provided a novel method to bypass the sterol exclusion barrier of aerobically grown yeast. Together with mass spectrometry (MS)-based lipidomics, we have found that MDM1 plays an active role in mediating sphingolipid metabolism. Collectively, our approach provides a new framework of analyzing lipid transporters without prior structural information or *in vitro* reconstitution.

## ACKNOWLEDGEMENTS

This work was supported by funds from the Welch Foundation (I-1873), the NIH NIGMS (GM119768), the NIDDK (R01DK126887), the Ara Parseghian Medical Research Fund, and the UT Southwestern Endowed Scholars Program (W.M.H.), and by the Leducq Foundation, the NCCR Chemical Biology and Swiss National Science Foundation (51NF40-185898 and 310030_184949) and the Canton of Geneva (H.R.).

## AUTHOR CONTRIBUTIONS

S.F., V.G. and H.R. conceived the idea and designed the experiments. S.F. synthesized caged compounds. H.H. and W.M.H. created the plasmids. V.G. and S.F. performed experiments and analyzed results. W.M.H and H.R. supervised the study. S.F. and H.R. wrote the manuscript with the help of all other authors.

---

**Supplementary Figure 1.**
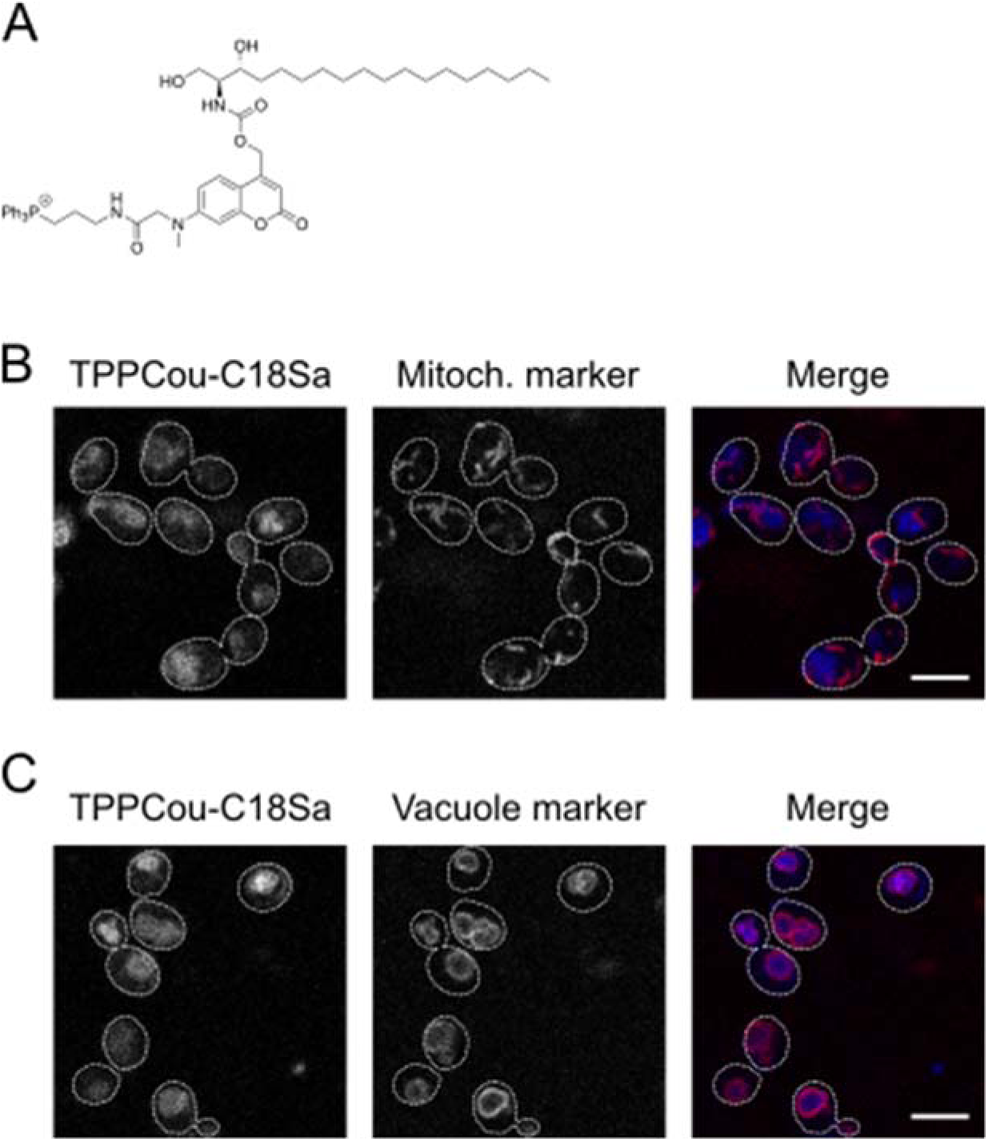
Mito-caged Sphinganine (Mito-Sa) probe is targeted to the vacuole in *Saccharomyces cerevisiae* (A) A chemical structure of TPPCou-C18Sa (B-C) Localization of TPPCou-C18Sa in yeast cells. Cells expressing mCherry-tagged mitochondrial marker Mdh1 (B) or vacuole marker Vph1 (C) were grown in rich-medium to log-phase were incubated with 2 µM TPPCou-C18Sa for 15 min at 30°C. Cells were washed, resuspended in low-fluorescence medium (LFM) and imaged as described. Cell outlines are shown by dashed lines. Bars, 5 µm.

**Supplementary Figure 2.**
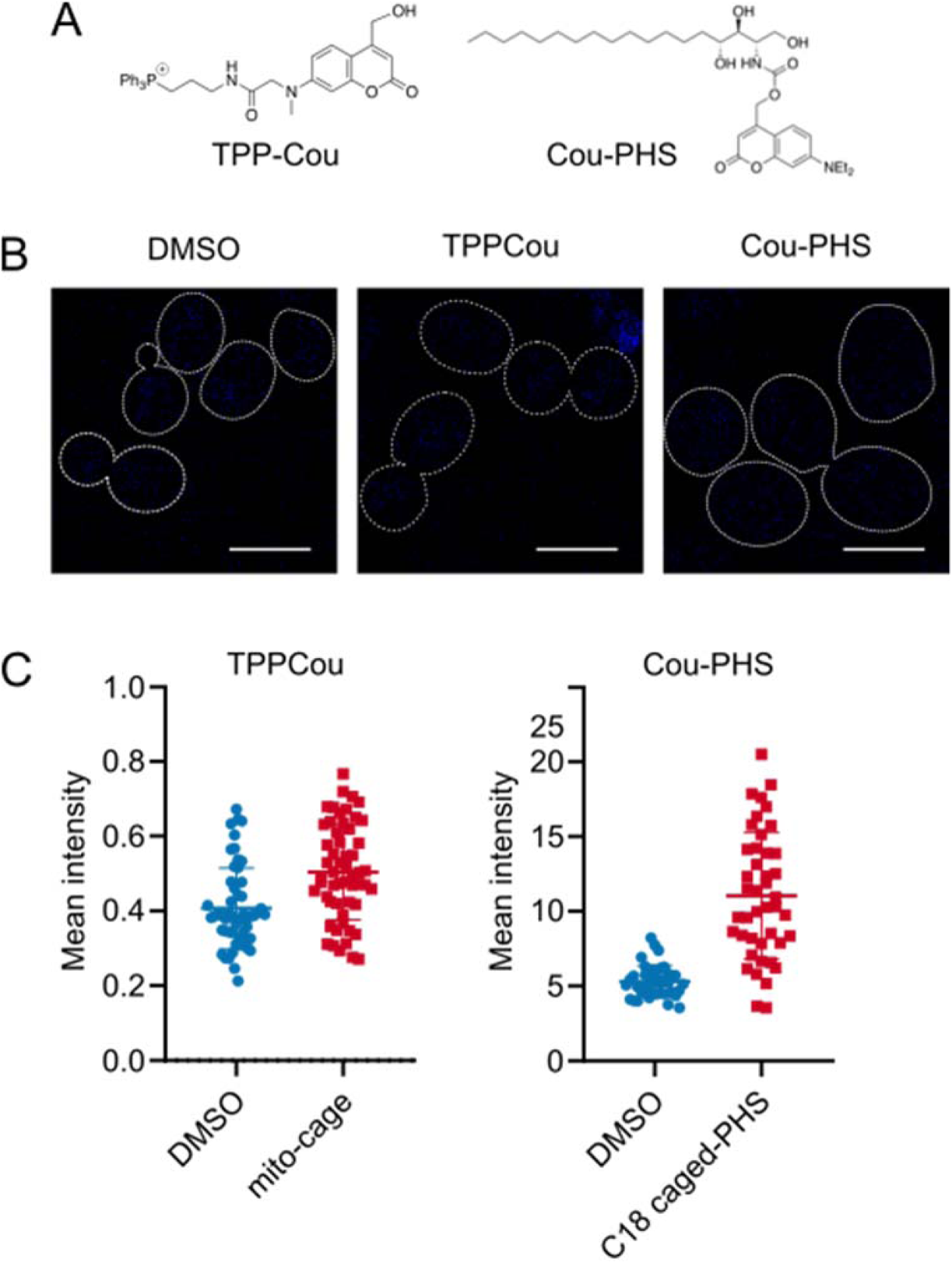
Caged PHS or mitochondrially-targeted cage alone are not internalized by the cells. (A) Chemical structures of Triphenylphosphonium-coupled coumarin (TPPCou) and diethylaminocoumarin caged PHS (Cou-PHS). (B) Representative fluorescence images of wild-type cells stained with 10 µM TPPCou or 50 µM Cou-PHS (both dissolved in DMSO) for 15 min at 30°C. Bars, 5 µm (C) Quantification of mean intracellular fluorescence intensity of coumarin from cells in (B). *n* > 50 cells, 3 biological replicates ns., not significant, student’s *t*-test.

**Supplementary Figure 3.**
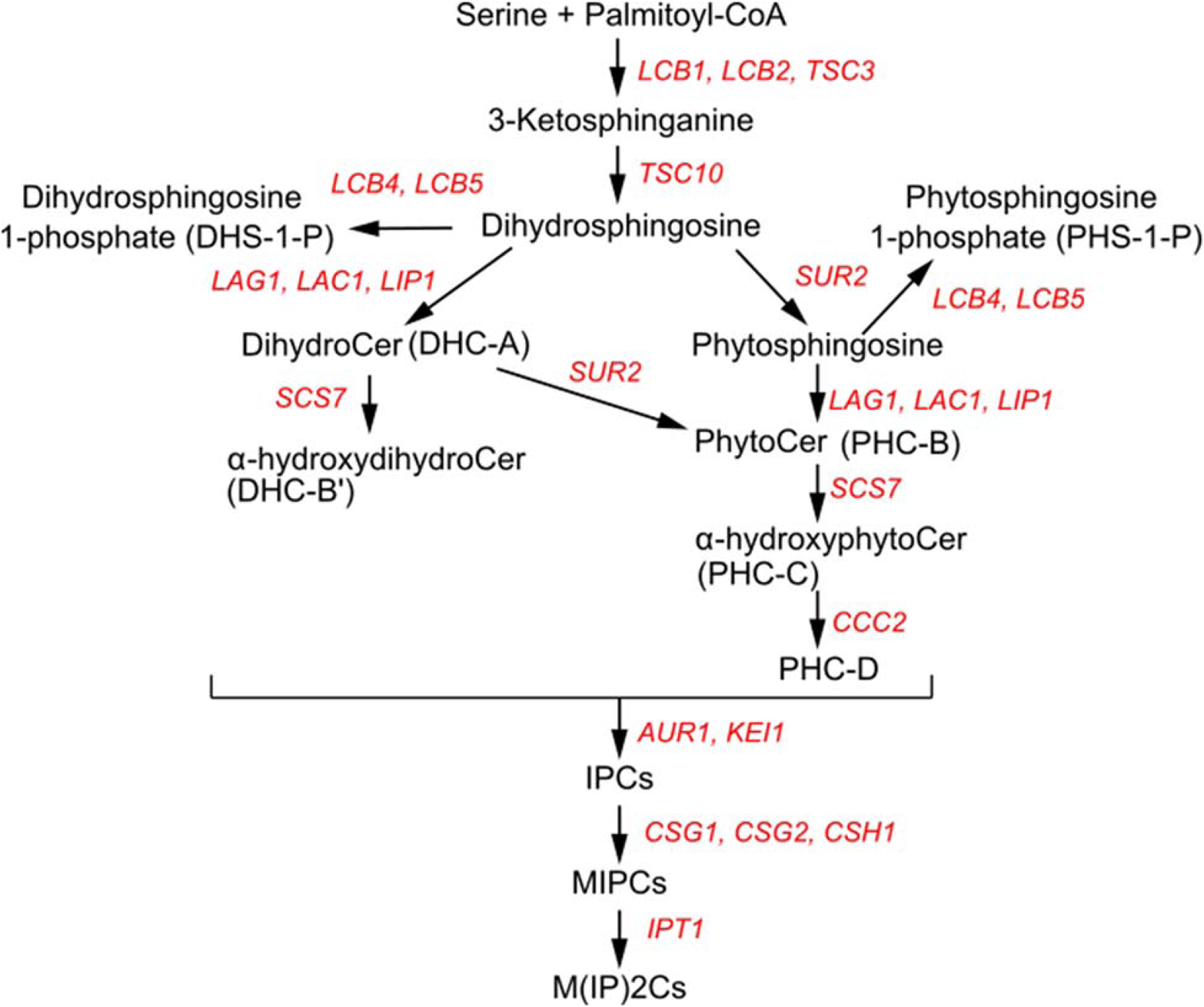
A simplified sphingolipid biosynthetic pathway in yeast. Lipids are shown in black and metabolic enzymes are shown in red color.

**Supplementary Figure 4.**
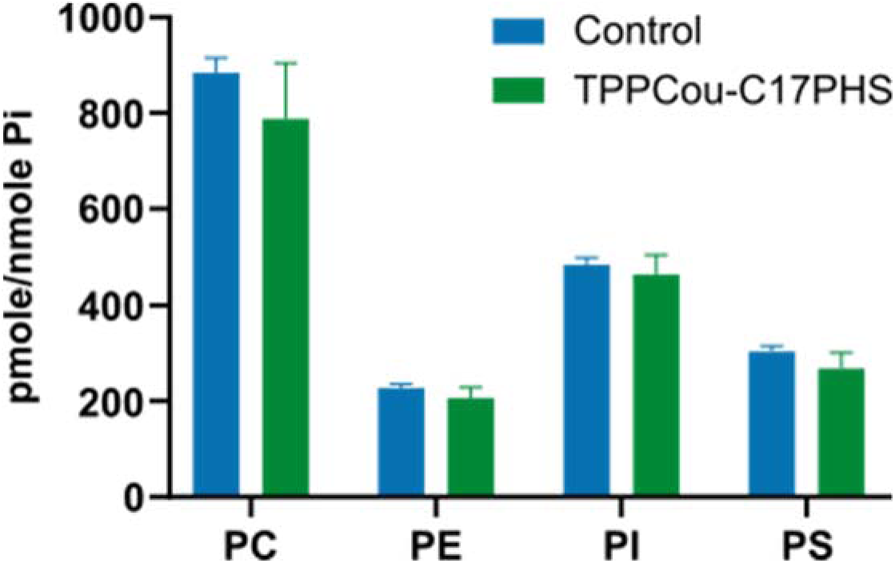
Glycerophospholipid profile of cells incubated with DMSO (Control) or 5 µM TPPCou-C17PHS. Data is normalized with respect to the total phosphate content. *n* = 3 biological replicates Error bars represent *SD*.

## Material and Methods

### Chemicals and reagents

All internal lipid standards, C17PHS (Phytosphingosine), C17Sa (Sphinganine) were purchased from Avanti Polar Lipids (Alabaster, AL). Yeast media were prepared from the following reagents: Glucose monohydrate, Bacto Peptone, Bacto yeast extract, Bacto agar were purchased from BD Biosciences (Sparks, MD); Yeast synthetic drop-out amino acid supplements, uracil and adenine were procured from Sigma (St. Louis, MO). Plasmids pBP73G-Mdm1-GFP and pBP73G-Nvj1-GFP were created as previously described ^1^.

### Yeast strains and culture

Baker’s yeast *Saccharomyces cerevisiae* were grown in standard rich medium supplemented with adenine and uracil (YPUAD −2% glucose, 1% yeast extract, 2% Bacto™ Peptone 2%, 10 mM MES (2-(N-morpholino)ethanesulfonic acid), 2% agar, 40 mg/l adenine and uracil, pH = 6.0) or in Synthetic Defined medium (SD −2% glucose, 0.67% Yeast Nitrogen Base (without amino acids, with ammonium sulfate), 1.92 g/l amino acid supplements, 2% agar, Na_2_HPO_4_ 10 mM, KH_2_PO_4_ 40 mM, pH = 6.0). Unless indicated otherwise, yeast cells were cultured at 30°C with vigorous shaking at 220 rpm. Yeast strains were constructed using standard techniques based on homologous recombination of PCR-generated linear fragments ^2^. Yeast strains used in the present study are listed in the Supplementary Table 1.

**Supplementary Table 1:**
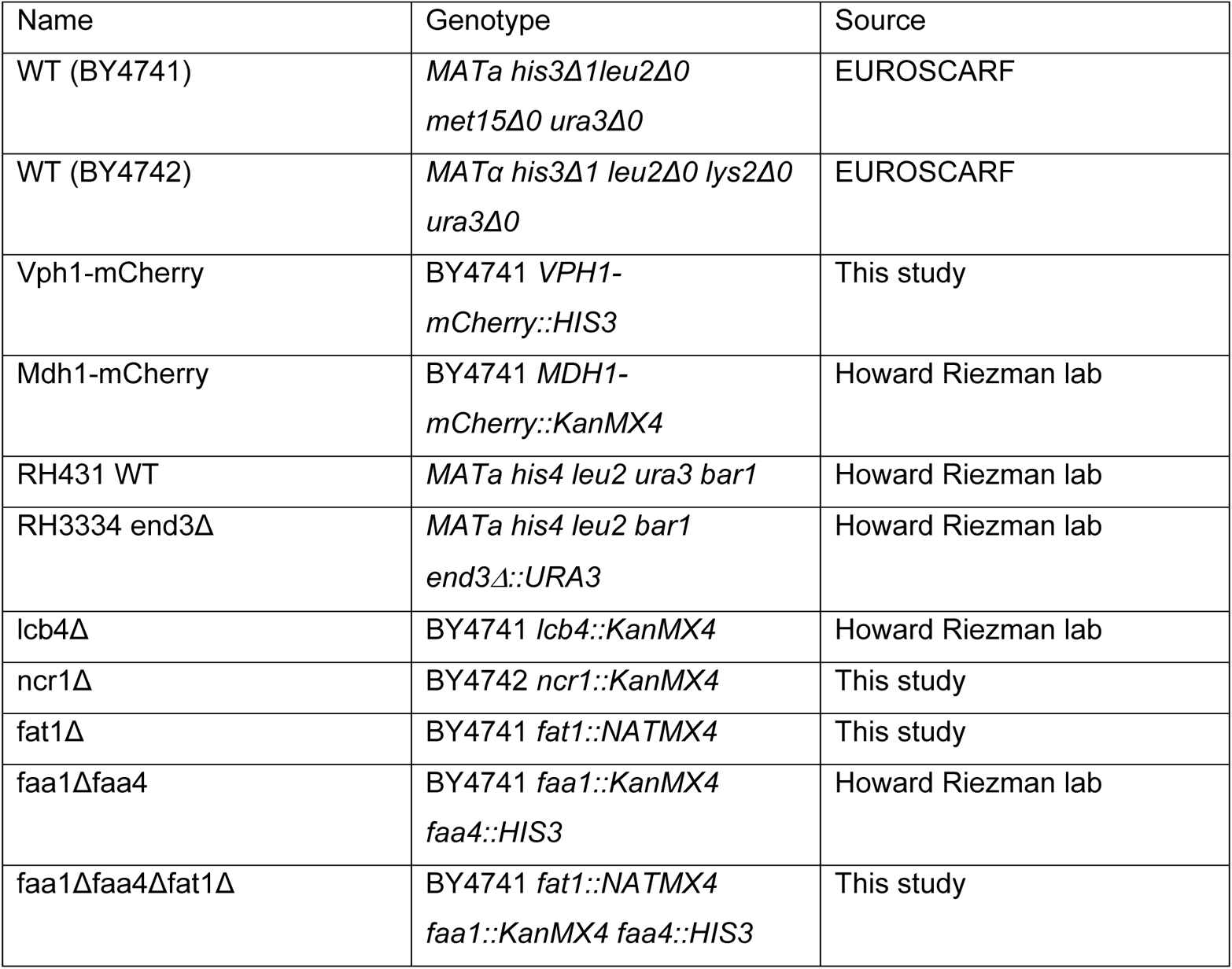
List of yeast strains used in this study

| Name | Genotype | Source |
| --- | --- | --- |
| WT (BY4741) | *MATa his3Δ1leu2Δ0*<br>*met15Δ0 ura3Δ0* | EUROSCARF |
| WT (BY4742) | *MATa his3Δ1 leu2Δ0 lys2Δ0*<br>*ura3Δ0* | EUROSCARF |
| Vph1-mCherry | BY4741 *VPH1-*<br>*mCherry::HIS3* | This study |
| Mdh1-mCherry | BY4741 *MDH1-*<br>*mCherry::KanMX4* | Howard Riezman lab |
| RH431 WT | *MATa his4 leu2 ura3 bar1* | Howard Riezman lab |
| RH3334 end3Δ | *MATa his4 leu2 bar1*<br>*end3Δ::URA3* | Howard Riezman lab |
| lcb4Δ | BY4741 *lcb4::KanMX4* | Howard Riezman lab |
| ncr1Δ | BY4742 *ncr1::KanMX4* | This study |
| fat1Δ | BY4741 *fat1::NATMX4* | This study |
| faa1Δfaa4 | BY4741 *faa1::KanMX4*<br>*faa4::HIS3* | Howard Riezman lab |
| faa1Δfaa4Δfat1Δ | BY4741 *fat1::NATMX4*<br>*faa1::KanMX4 faa4::HIS3* | This study |

### Yeast growth monitoring

Yeast cells were grown overnight in liquid YPUAD or YPUAG were diluted to OD600 = 0.1 in the same medium, 200 μl of cells suspensions were put into flat-bottom 96-well plates. The plate was sealed with gas permeable moisture barrier seal (4titude) and absorbance at a wavelength of 600 nm (OD600) was recorded every 10 minutes over 25 hours at 30℃using a plate reader (Biotek™ Synergy™ H1, BioTec Instruments, Winooski, VT).

### Fluorescence microscopy and image analysis

Cells were grown in YPUAD or selective SD medium to the exponential phase, washed and resuspended in low-fluorescence medium (LFM) – SD medium lacking folate and riboflavin. 15 μl of cells suspension were applied onto Concanavalin-A pre-treated glass slides and incubated at room temperature for 5 min. The unattached cells were gently aspirated, the slides were covered with a coverslip and sealed with nail polish. The samples were imaged with LSM700 Zeiss confocal microscope using a 100x oil immersion objective. Quantification of fluorescence intensity was carried out using ImageJ software. Pearson’s correlation coefficients were calculated using EzColocalization ImageJ plugin ^3^.

### Labeling with synthetic chemical probes

Yeast cells were grown in YPUAD or SD medium to the logarithmic phase, labeled with TPP-Cou probes for 20 minutes at 30℃, centrifuged at 800g, resuspended in LFM medium and processed for imaging as described above.

### Photo-uncaging experiments

UV uncaging experiments were carried-out using a 1000-Watt Arc Lamp Source (#66924, NewPort) equipped with a dichromic mirror (350–450 nm, #66226). Yeast cells were grown in YPUAD at 30℃with shaking (220 rpm) to mid-logarithmic phase corresponding to OD600 value of 1.2-1.5. Cells loaded with TPPCou-caged compounds by incubation at 30 °C. After incubation with the caged compounds, cells were placed on ice, washed with YPUAD medium and subjected to UV uncaging. Cell were placed on ice at a distance of 20 cm from the UV-lamp and were irradiated for 120 s at 1000 Watt. In time-course experiments cells were re-suspended in warm YPUAD medium after UV-uncaging and were further cultured at 30℃ for the indicated time periods.

### LC-MS/MS measurement of sphingoid bases^6^

Yeast cells grown at 30°C in YPUAD medium to exponential phase (OD600 ∼ 1.2-1.5) were labelled with TPPCou-caged compounds and subjected to UV illumination. Irradiated cells were pelleted, washed in 5 ml ice-cold water, resuspended in 1 ml cold water and the amount corresponding to 5 OD_600_ units were collected for measurement. To extract metabolites, 150 µl of extraction solvent (water, ethanol, diethyl ether, pyridine, 4.2 N ammonium hydroxide 15:15:5:1:0.018, vol/vol) together with a mix of internal standards (0.04 nmol of C17 sphinganine, 0.4 nmol of C17 sphinganine-1-phosphate) and 100 µl of Zirconium oxide beads (Bertin Technologies, Montigny-le-Bretonneux, France) were added to cell pellets. Cells were broken using a cryolysis breaker (Bertin Technologies) at 0°C, incubated for 20 min on ice and centrifuged at 14000 rpm for 2 min at 0 °C. Supernatants were collected and 150 µl of extraction solvent were added again followed by brief vortexing and centrifugation. The supernatants from the two extraction steps were combined and centrifuged at 14000 rpm for 2 min. The extracts were vacuum-dried in CentriVap (Labconco). To the lipid film, a mixture consisting of 70 µl of borate buffer (200 mM boric acid pH 8.8, 10 mM tris(2-carboxyethyl)-phosphine, 10 mM ascorbic acid and 33.7 mM 15N13C-valine), and 10 ml of formic acid solution (0.1%in water) was added for derivatization by the reaction with 20 µl AQC (6-aminoquinolyl-N-hydroxysuccinimidyl carbamate) solution (2.85 mg/ml in acetonitrile) at 55°C for 15 min. After that the samples were incubated overnight at 24°C. AQC-derivatized samples were analyzed by LC-MS/MS using an Accela-HPLC system coupled with a TSQ Vantage Triple Quadrupole mass spectrometer (Thermo Fisher Scientific). Selected metabolites were detected in a multiple reaction monitoring (MRM) setup detecting the AQC fragment. Signals were normalized with respect to the values of ^15^N^13^C-valine and the corresponding internal standards.

### Lipidomics

#### Sample collection and lipid extraction

Cells grown to exponential phase (OD600 ∼ 1.2-1.5) were loaded with 5 µM TPP-Cou-C17PHS or DMSO (vehicle) for 20 min at 30°C, washed with cold YPUAD medium followed by UV-uncaging on ice for 2 min as described. After uncaging TPP-Cou-C17PHS-treated cells were resuspended in fresh medium and were cultured for 20, 40 or 60 min followed by treatment with 5% trichloracetic acid (TCA) for sample collection. The extraction procedure was performed as described in ^4^ with minor modifications. First, a mix containing internal lipid standards (7.5 nmol 17:0/14:1 PC, 4.0 nmol 17:0/14:1 PS, 6.0 nmol 17:0/14:1 PI 7.5 nmol 17:0/14:1 PE, 1.2 nmol C17 ceramide and 2.0 nmol C8 glucosylceramide (C8GC)) was added to the cell pellets together with 1.5 ml of extraction solvent (H_2_O, ethanol, diethyl ether, pyridine, 4.2 N NH_4_OH 15:15:5:1:0.018) and 500 µl glass beads. Cells were disrupted by vigorous vortexing on a multivortexer for 6 minutes at room temperature, followed by the incubation at 60°C for 20 minutes. After centrifugation at 800 g, the supernatant was transferred to fresh 13 mm glass tubes. Then, the extraction procedure was repeated. Supernatants from the two rounds of extraction were combined and split into 2 equal aliquots: for glycerophospholipid and sphingolipid analysis. The extracts were dried either under the flow of N_2_ at 80°C or vacuum-dried in a CentriVap, flushed with nitrogen flow and stored at −80°C until usedfor further steps. The fractions for sphingolipid analysis were subjected to mild alkaline hydrolysis to deacylate glycerophospholipids that are a source of ion suppression when using direct infusion techniques. For this the dried samples were re-suspended by sonication in 1 ml monomethylamine reagent (methanol, H_2_O, n-butanol, methylamine 4:3:1:5 vol/vol/vol), incubated at 53°C for 1 hour, dried and stored at −80°C. Finally, sample desalting was accomplished by n-butanol extraction. Dried samples were re-suspended in 300 µl water-saturated n-butanol followed by addition of 150 µl of LC-MS-grade water. In order to induce phase separation, the samples were centrifuged at 3200 g for 10 minutes at 20°C, the upper phases were collected. The desalting procedure was repeated twice, the combined extracts were dried and frozen at −80°C.

#### Direct-Infusion Lipidomics^5^

Dried samples were resuspended by sonication in 500 µl of LC-MS-grade chloroform:methanol (1:1, vol/vol) followed by dilution in chloroform:methanol:water (2:7:1, vol/vol/vol) or in chloroform:methanol (1:2, vol/vol) containing 5 mM ammonium acetate for positive or negative mode MS analysis, respectively. The samples containing sphingolipids were diluted 1:20 (vol/vol). Lipid mixtures were transferred into wells of a 96-well plate which was sealed with an aluminum foil before MS analysis. Samples were injected into the TSQ Vantage Triple Quadrupole mass spectrometer (Thermo Fisher Scientific, Waltham, MA) equipped with the TriVersa Nanomate - automated chip-based electrospray ionization platform (Advion Bioscience, Ithaca, NY). Samples were injected at a gas pressure of 30 psi with a spray voltage of 1.2 kV and run on the mass spectrometer operated with a spray voltage of 3.5 kV for positive mode and 3.0 kV for negative mode and a capillary temperature set at 190°C. Sphingolipid species were identified and quantified using multiple reaction monitoring (MRM). Each lipid species was quantified using standard curves created from internal standards. Data analysis was performed using R language.

#### Reversed-phase UHPLC-HRMS analyses

Dried samples were resuspended by sonicating in 100 µl of LC-MS-grade chloroform:methanol (1:1, vol/vol). Reversed-phase UHPLC-HRMS analyses were performed using a Q Exactive Plus Hybrid Quadrupole-Orbitrap mass spectrometer coupled to an UltiMate 3000UHPLC system (Thermo Fisher Scientific) equipped with an Accucore C30 column (150 x 2.1 mm, 2.6 μm) and its 20mm guard (Thermo Fisher Scientific). Samples were kept at 8°C in the autosampler, 10 μl were injected and eluted with a gradient starting at 10% B for 1 min, 10-70% B in 4 min, 70-100% B in 10 min, washed in 100 % B for 5 min and column equilibration for an additional 3 min. Eluents were made of 5 mM ammonium acetate and 0.1% formic acid in water (*solvent A*) or in isopropanol/acetonitrile (2:1, v/v) (*solvent B*). Flow rate and column oven temperature were respectively at 350μl/min and 40°C. The mass spectrometer was operated using a heated electrospray-ionization (HESI) source in positive and negative polarity with the following settings: electrospray voltage: −3.4 KV (−) or 3.9 KV (+); sheath gas: 51; auxiliary gas: 13; sweep gas: 3; vaporizer temperature: 431°C; ion transfer capillary temperature: 320 °C; S-lens: 50; resolution: 140,000; m/z range: 200-1000; automatic gain control: 1e6; maximum injection time: 50 ms. For identification of phytoceramides and IPCs, parallel reaction monitoring (PRM) measurement was performed using a predetermined inclusion list of corresponding lipid species. The following setting was used in HCD fragmentation: automatic gain control: 2.5e5; maximum injection time: 120 ms; resolution: 35,000; (N)CE: 30. Xcaliburv.4.2 (Thermo Fisher Scientific) was used for data acquisition and processing.

### Total phosphate measurement

Total lipid extracts were re-suspended in 500 µl of LC-MS-grade chloroform:methanol (1:1, vol/vol), 50 µl were transferred into Pyrex tubes and dried. Different amounts of standard solution of 5 mM KH_2_PO_4_ were prepared for the standard curve. 140 µl of perchloric acid (70%) and 20 µl of water were added to the dried lipids and standards and the tubes were incubated in a heating-block at 100°C for 1h in a fume hood. Tubes were cooled for 5 min at RT and 800 µl of freshly-prepared reagent solution (water:1.25% ammonium molybdate:1.67% ascorbic acid, 5:2:1, vol/vol) was added and the samples were heated at 180°C for 5 min. After cooling at RT, 100 µl was used to measure absorbance at 820 nm.

### Chemical synthesis

Chemicals and reagents were purchased from commercial sources and were used without further purification unless otherwise mentioned. d7-Sphinganine and C17PHS were purchased from Avanti Polar Lipids. Deuterated solvents were obtained from Cambridge Isotope Laboratories, Inc. ^1^H and ^13^C-NMR spectra were recorded on a Bruker AMX-400 MHz or 500 MHz spectrometer. Chemical shifts are given in ppm (δ) using the NMR solvent as internal references and J values are reported in Hz. Splitting patterns are designated as follows: s, singlet; d, doublet; t, triplet; q, quartet; m, multiplet; b, broad. ^13^C-NMR spectra were broadband hydrogen decoupled. LC-MS were recorded using a Thermo Electron Corporation HPLC with a Thermo Scientific Finnigan Surveyor MSQ Spectrometer System. Reverse phase HPLC purification was performed using an Agilent Technologies 1260 infinity HPLC equipped with a ZORBAX 300 SB-C18 column (9.4 x 250 mm). High-resolution mass spectra were recorded on a Thermo Q Exactive Plus mass spectrometer. Thin layer chromatography (TLC) was performed on aluminum-backed, pre-coated silica gel plates (Merck TLC silica gel 60 F254). Spots were detected by a UV lamp under 254 nm or 365 nm wavelength. All reactions were carried out under nitrogen atmosphere unless otherwise mentioned.

### TPPCou-C17PHS

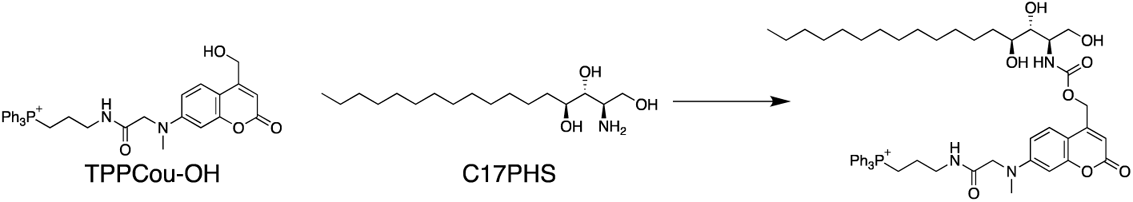

TPPCou-C17PHS was synthesized according to our previously described procedures. To a solution of TPP-Cou-OH (26 mg, 0.04 nmol) in 4 mL of dry DMF, Bis-(4-nitrophenyl)carbonate (12 mg, 0.04 mmol) and DIPEA (20 μL, 0.11 mmol) was added. The reaction mixture was stirred at room temperature in dark. After 3 hr, C17PHS (11 mg, 0.04 mmol) in 1 mL of DMF was added and the reaction was stirred at 60 °C for 3 hr in dark. The crude product was concentrated under reduced pressure and purified by flash chromatography (SiO_2_, MeOH/DCM = 1/10) to afford TPPCou-C17PHS (18 mg, Yield: 45 %) as a yellow solid. The product was further purified by reverse phase HPLC. Semi-preparative HPLC runs were carried out with a gradient from 5 % to 95 % acetonitrile/water system (0.1 % TFA) for 20 min and a flow of 1 mL/min, monitored by a PDA detector at 254 nm and 360 nm. The fraction was collected and lyophilized to afford 4.85 mg of the product. ^1^H NMR (400 MHz, DMSO-d_6_) δ = 8.17 (t, *J* = 5.7 Hz, 1H), 7.88 (m, 3H), 7.70-7.77 (m, 12H), 7.42 (d, *J* = 8.9 Hz, 1H), 7.13 (d, *J* = 8.9 Hz, 1H), 6.65 (dd, *J* = 8.9, 2.2 Hz, 1H), 6.51 (d, *J* = 2.2 Hz, 1H), 6.60 (s, 1H), 5.17 (m, 2H), 4.40 (s, 2H), 3.75 (t, *J* = 5.0 Hz, 1H), 3.60-3.71 (m, 2H), 3.43-3.52 (m, 3H), 3.37 (m, 2H), 3.25 (m, 2H), 3.07 (s, 2H), 1.66 (m, 2H), 1.39-1.51 (m, 2H), 1.18-1.25 (m, 24H), 0.83 (t, *J* = 6.9 Hz, 3H) ppm. ^13^C NMR (101 MHz, DMSO-d_6_) δ = 134.93, 134.91, 133.49, 133.39, 130.26, 130.14, 124.79, 109.07, 105.08, 97.68, 74.61, 70.72, 63.25, 60.58, 60.23, 54.95, 54.44, 42.74, 38.67, 38.48, 31.94, 31.24, 29.15, 29.06, 29.02, 28.96, 28.66, 25.25, 22.21, 22.04, 18.53, 18.01, 13.89 ppm. HR-ESI-MS (pos.): C_52_H_69_N_3_O_8_P, [M]^+^ calculated: 894.48223, found: 894.48104.

### TPPCou-Sa_D7

The synthetic procedures of TPPCou-Sa (previously named as Mito-Sa) have been reported ^6^. The synthesis of stable isotope-labeled TPPCou-Sa_D7 was carried out in smaller scale and 0.2 mg of the final compound was obtained after HPLC purification. Due to the low amount, this compound was only characterized by LC-MS and HR-MS. HR-ESI-MS (pos.): C_52_H_62_D_7_N_3_O_7_P, [M]^+^ calculated: 899.5469, found: 899.54559.

### TPPCou-Chol

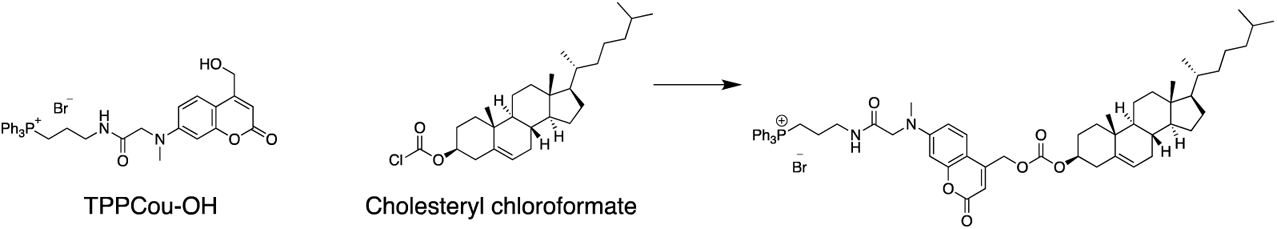

To a solution of TPPCou-OH ^6^ (bromide salt, 64 mg, 0.1 mmol) in 5 mL dry DCM, cholesteryl chloroformate (43 mg, 0.1mmol, 1.0 equiv.) and DMAP (13 mg, 0.11 mmol, 1.1 equiv.) were added and the reaction mixture was allowed to stir at room temperature for 18 h in dark. The crude product was extracted by DCM/H_2_O, washed by brine, concentrated under reduced pressure, and purified by flash chromatograph (SiO_2_, 5-10 % MeOH in DCM) to afford TPPCou-Chol as a yellow solid (60 mg, Yield: 60 %). ^1^H NMR (400 MHz, CDCl_3_) δ = 9.31 (s, 1H), 7.75 (m, 3H), 7.60-7.69 (m, 12H), 7.14 (d, *J* = 8.9 Hz, 1H), 6.77 (dd, *J* = 8.9, 1.9 Hz, 1H), 6.44 (d, *J* = 1.9 Hz, 1H), 6.00 (s, 1H), 5.38 (d, *J* = 5.0 Hz, 1H), 5.07 (s, 2H), 4.48 (m, 1H), 4.21 (s, 2H), 3.56-3.61 (m, 2H), 3.48 (s, 2H), 3.16 (s, 3H), 2.62 (bs, 2H), 2.38 (m, 2H), 2.13 (s, 2H), 1.83-1.92 (m, 6H), 0.99-1.67 (m, 24H), 0.89 (d, *J* = 6.5 Hz, 3H), 0.83 (d, *J* = 1.7 Hz, 3H), 0.81 (d, *J* = 1.7 Hz, 3H), 0.64 (s, 3H) ppm. ^13^C NMR (101 MHz, CDCl_3_) δ = 170.21, 161.45, 155.65, 153.95, 152.66, 149.00, 139.12, 135.20, 135.17, 133.44, 133.34, 130.63, 130.51, 124.07, 123.16, 118.45, 117.59, 109.83, 106.65, 106.50, 98.48, 78.77, 64.07, 56.66, 56.11, 55.87, 49.97, 42.29, 40.31, 39.69, 39.49, 38.88, 38.71, 37.95, 36.80, 36.52, 36.16, 35.76, 31.89, 31.81, 30.92, 29.61, 28.20, 27.99, 27.64, 24.26, 23.80, 22.81, 22.55, 22.47, 21.02, 20.41, 19.26, 18.70, 11.84 ppm. HR-ESI-MS (pos.): C_62_H_78_N_2_O_6_P, [M]^+^ calculated: 977.55975, found: 977.55862.

### Cou-C18PHS

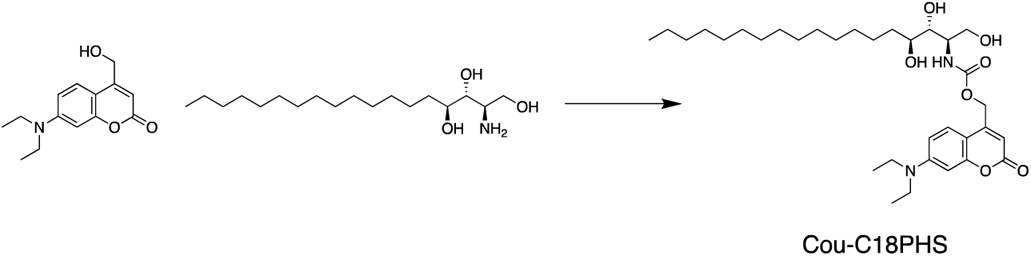

To a solution of 7-Diethylamino-4-hydroxymethylcoumarin ^7^ (25 mg, 0.1 mmol) and DIPEA (100 µL) in 3 mL of dry THF at 0 °C, triphosgene (30 mg, 0.1 mmol) dissolved in 1 mL of THF was added. The reaction mixture was stirred at 0 °C for 3 h before extracted by EtOAc/H_2_O. The organic phase was washed by brine, dried over Na_2_SO_4_, and concentrated under reduced pressure. The crude product, after re-dissolved in 2 mL dry THF, was slowly added into a solution of phytosphingosine (40 mg, 0.125 mmol, 1.25 equiv.) and DIPEA (50 µL) in 3 mL of dry THF via a syringe at 0 °C. The reaction mixture was kept in dark at room temperature for 1 h, and then was extracted by EtOAc/H_2_O, washed by saturated NaHCO_3_, brine, dried over Na_2_SO_4_, and concentrated under reduced pressure. The crude product was purified by flash chromatograph (SiO_2_, EtOAc/cyclo-hexane = 1/1, Rf ≈ 0.5), and further purified by a second flash chromatograph (SiO_2_, 5% MeOH in DCM, Rf = 0.2) to afford Cou-C18PHS as a yellow solid (57 mg, Yield: 96 %). ^1^H NMR (400 MHz, DMSO-d_6_) δ = 7.44 (d, *J* = 9.0 Hz, 1H), 7.13 (d, *J* = 9.0 Hz, 1H), 6.67 (dd, *J* = 9.0, 2.4 Hz, 1H), 6.52 (d, *J* = 2.4 Hz, 1H), 6.04 (s, 1H), 5.13-5.28 (m, 2H), 4.51-4.60 (m, 2H), 4.31 (d, *J* = 4.5 Hz, 1H), 3.61-3.67 (m, 2H), 3.17-3.43 (m, 7H, overlapped with water peak), 1.42-1.51 (m, 2H), 1.18-1.28 (m, 24H), 1.12 (t, *J* = 6.8 Hz, 6H), 0.84 (t, *J* = 6.8 Hz, 3H) ppm. ^13^C NMR (101 MHz, DMSO-d_6_) δ = 160.75, 155.72, 155.12, 151.97, 150.37, 125.2, 108.66, 105.25, 104.38, 96.83, 74.54, 70.71, 60.70, 60.42, 54.49, 43.98, 31.67, 31.31, 29.15, 29.11, 29.09, 29.03, 28.72, 25.30, 22.10, 13.95, 12.29 ppm. HR-ESI-MS (pos.): C_33_H_54_N_2_O_7_, [M+H]^+^ calculated: 591.40038, found: 591.40046.

---

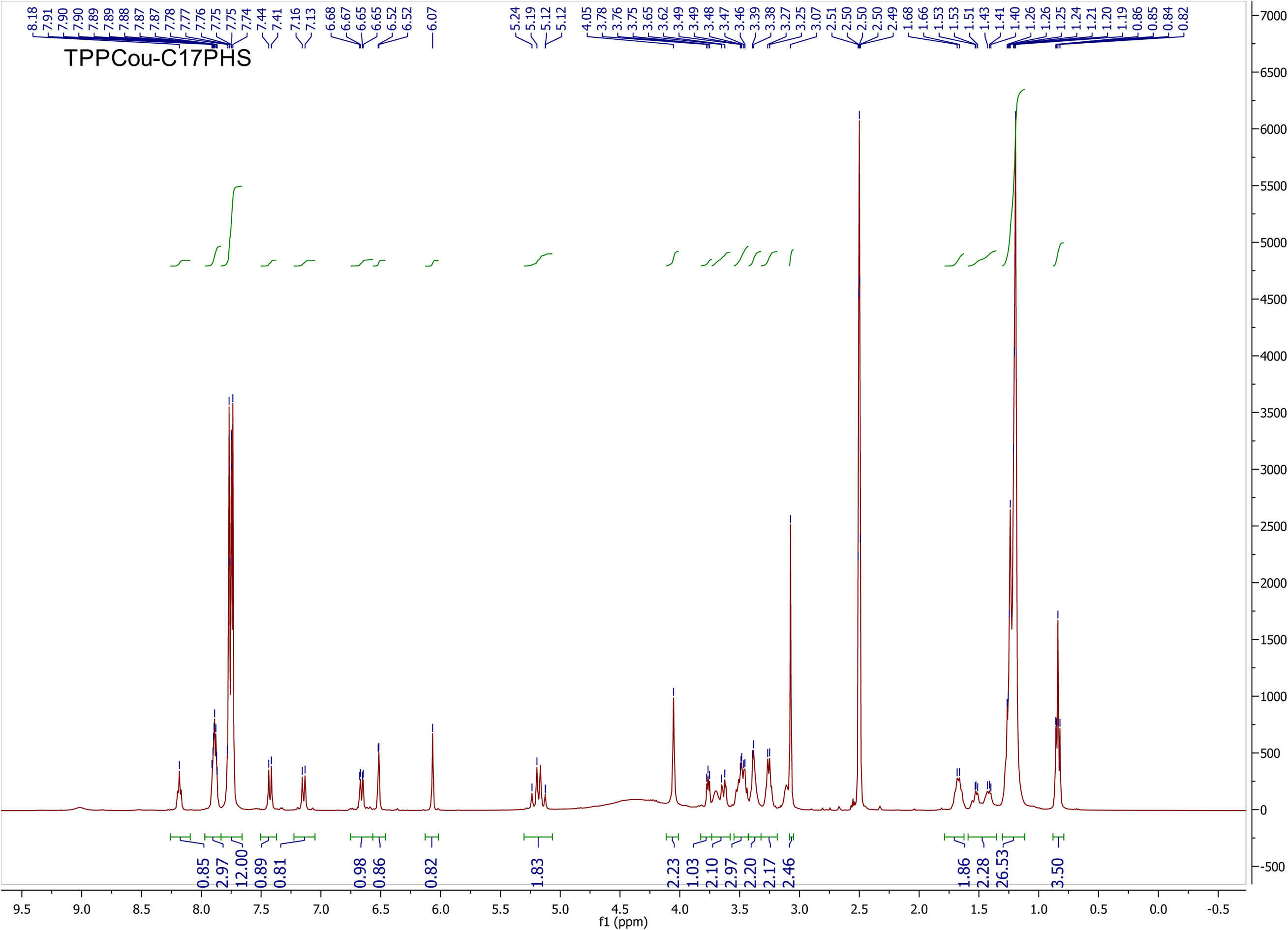

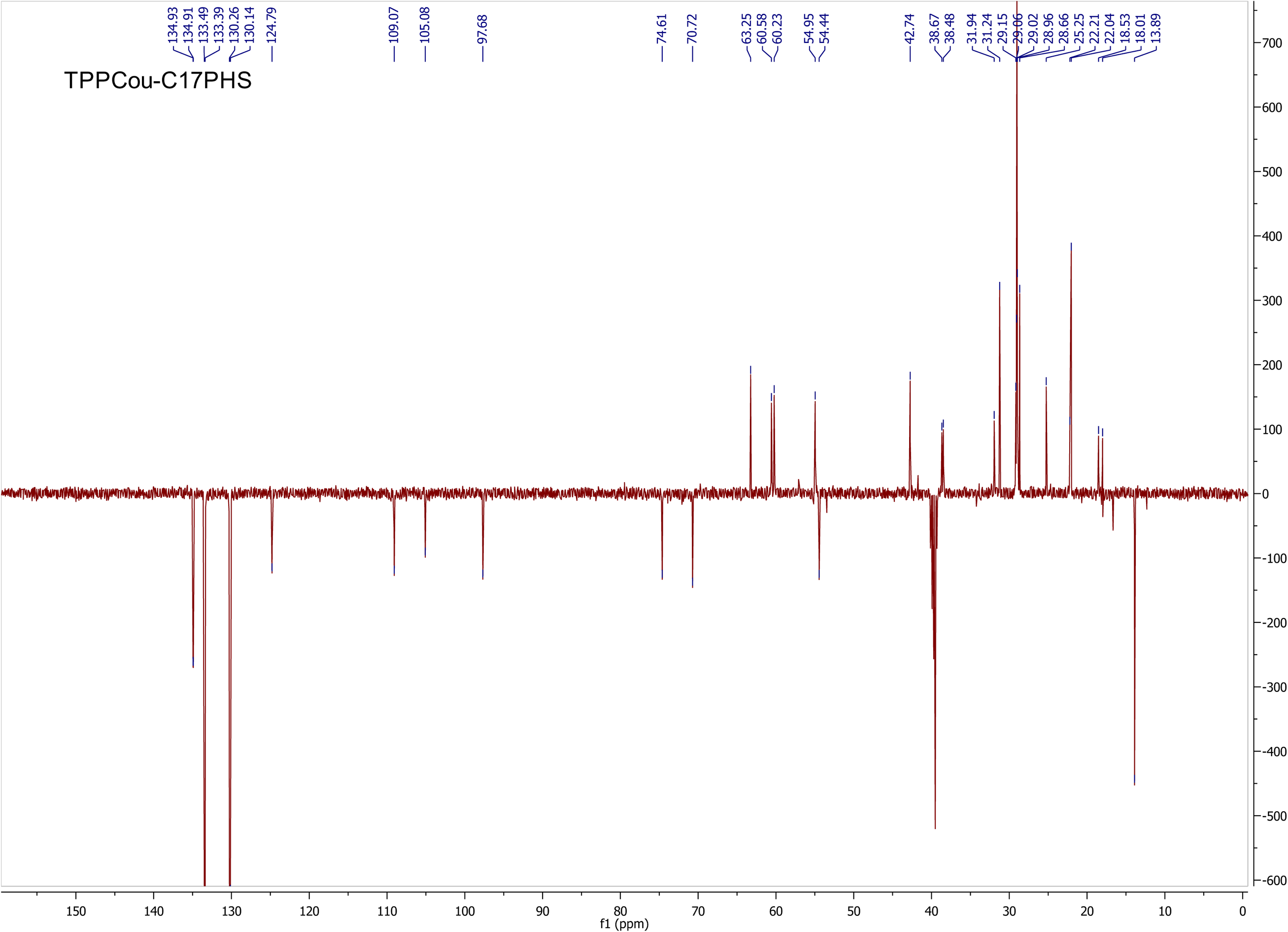

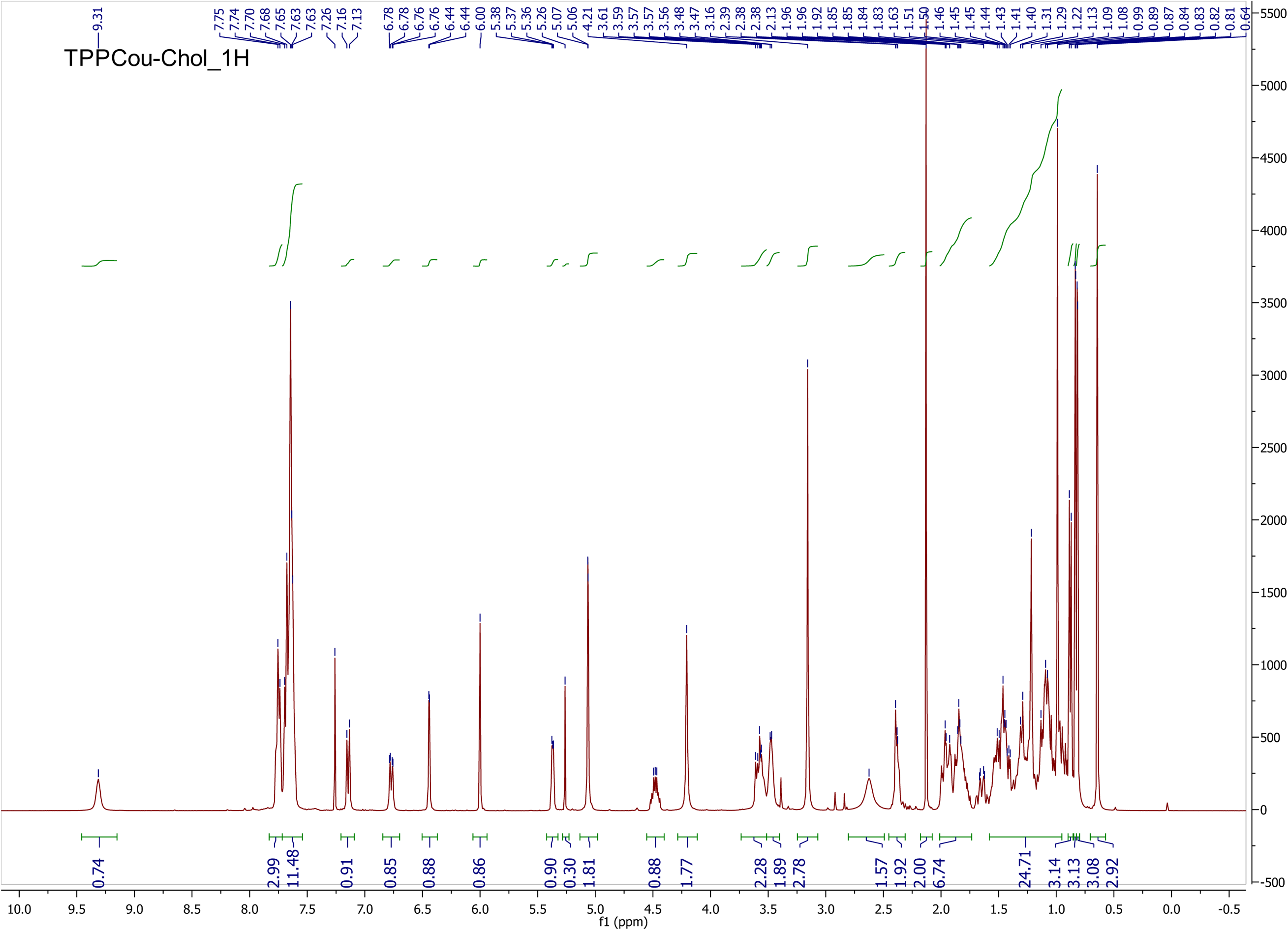

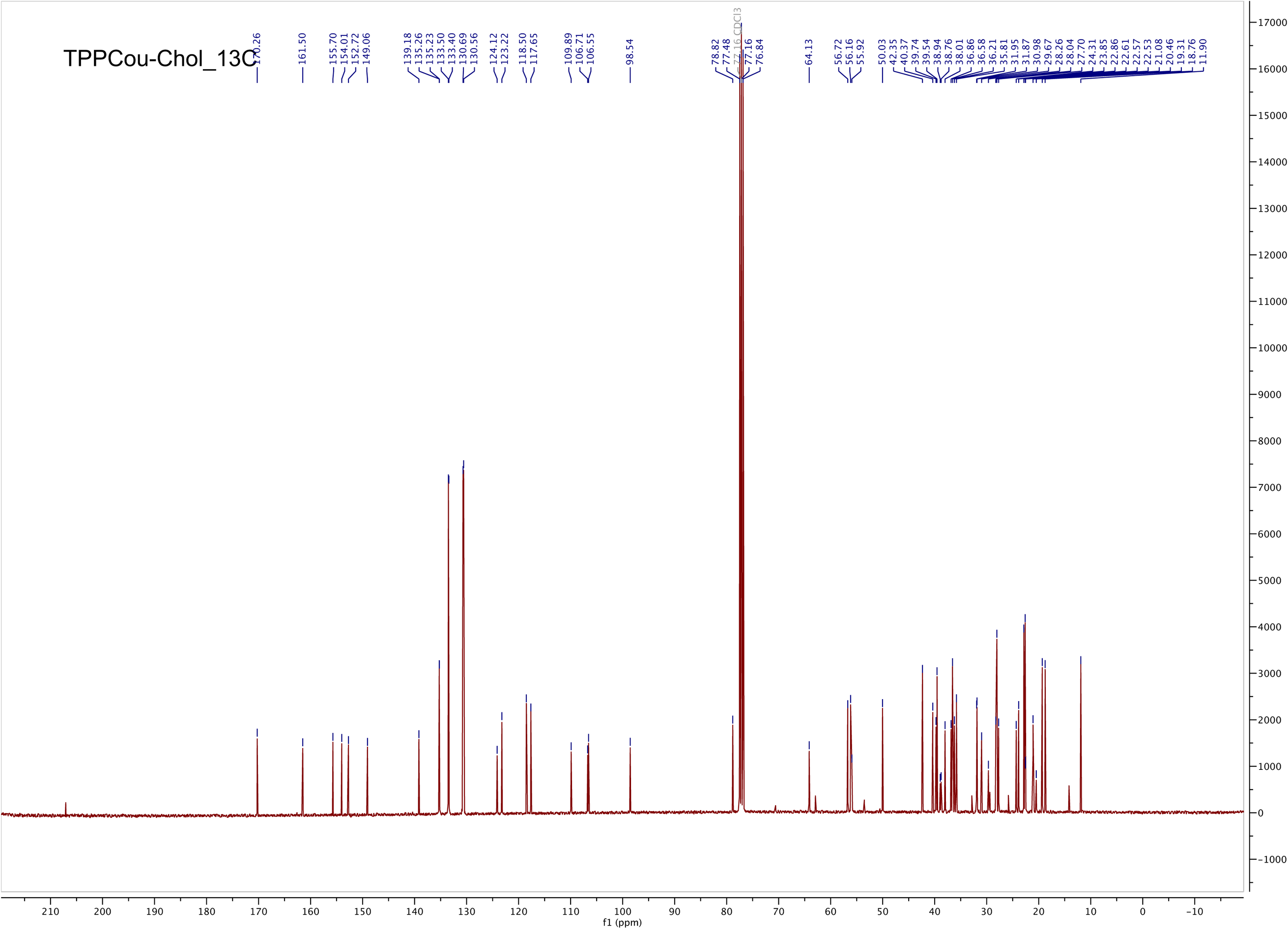

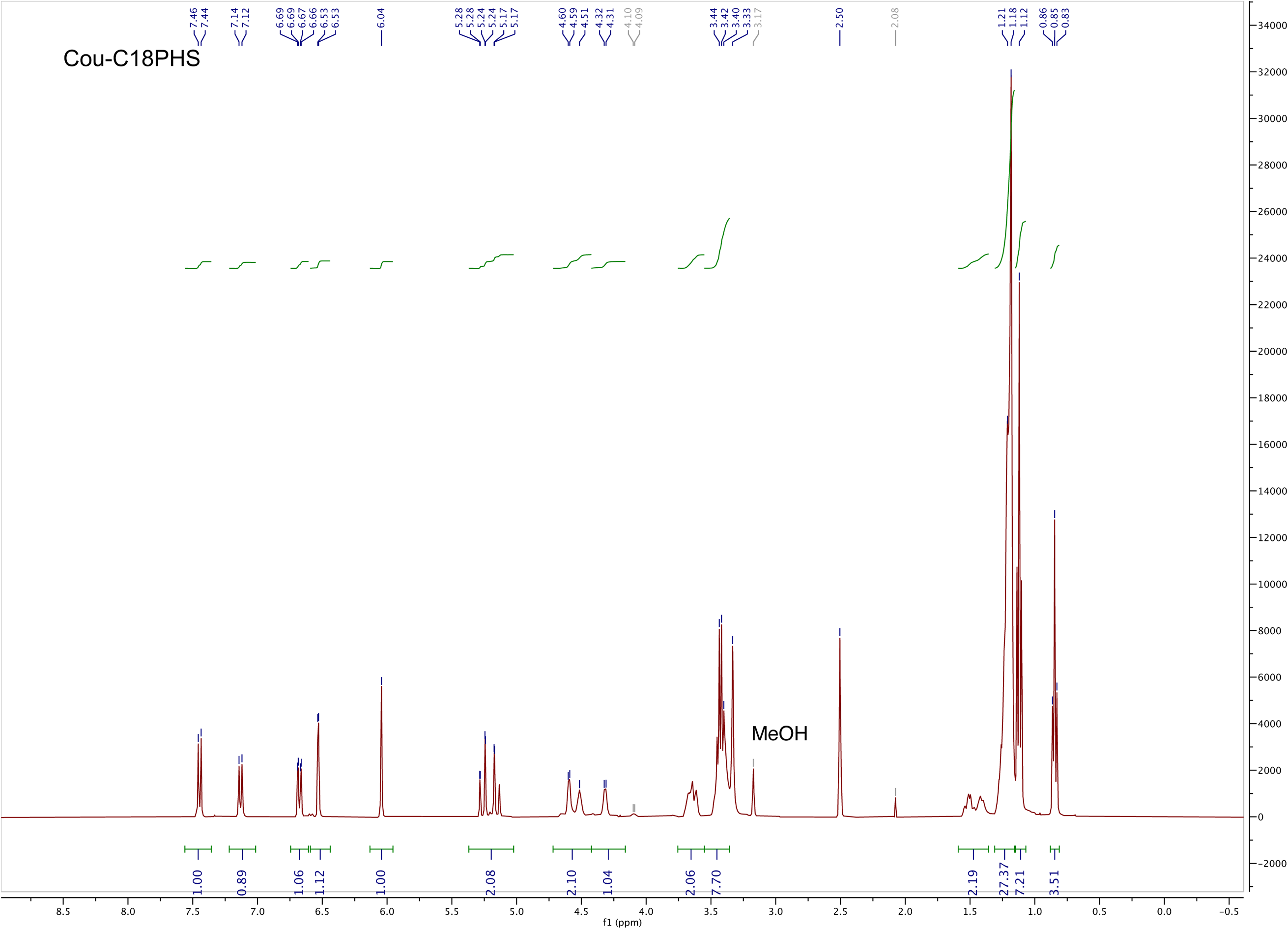

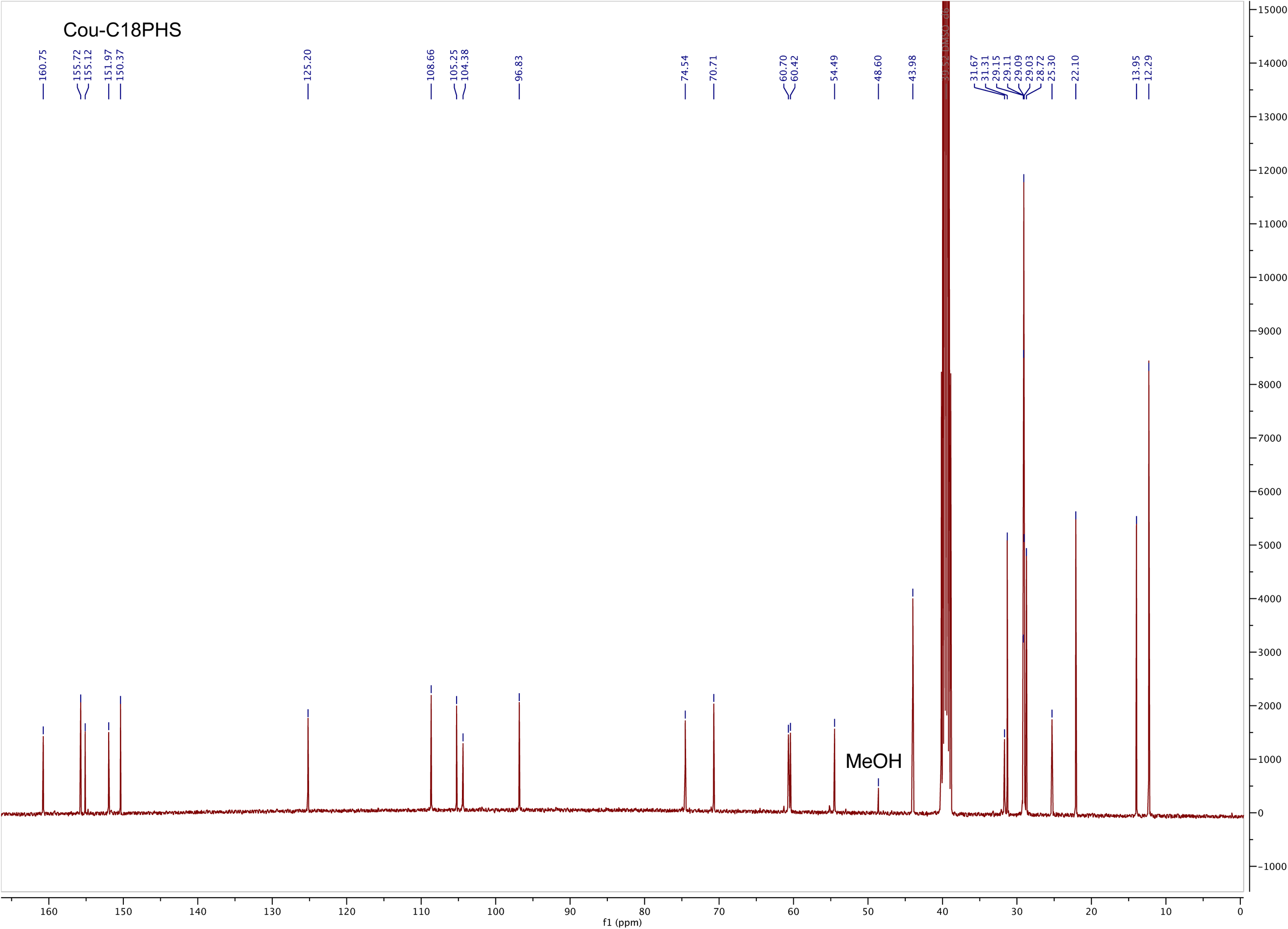

